# Principles of mRNA control by human PUM proteins elucidated from multi-modal experiments and integrative data analysis

**DOI:** 10.1101/2020.04.01.019836

**Authors:** Michael B. Wolfe, Trista L. Schagat, Michelle T. Paulsen, Brian Magnuson, Mats Ljungman, Daeyoon Park, Chi Zhang, Zachary T. Campbell, Aaron C. Goldstrohm, Peter L. Freddolino

## Abstract

The human PUF-family proteins, PUM1 and PUM2, post-transcriptionally regulate gene expression by binding to a PUM recognition element (PRE) in the 3’ UTR of target mRNAs. Hundreds of PUM1/2 targets have been identified from changes in steady state RNA levels; however, prior studies could not differentiate between the contributions of changes in transcription and RNA decay rates. We applied metabolic labeling to measure changes in RNA turnover in response to depletion of PUM1/2, showing that human PUM proteins regulate expression almost exclusively by changing RNA stability. We also applied an in vitro selection workflow to precisely identify the binding preferences of PUM1 and PUM2. By integrating our results with prior knowledge, we developed a ‘rulebook’ of key contextual features that differentiate functional vs. non-functional PREs, allowing us to train machine learning models that accurately predict the functional regulation of RNA targets by the human PUM proteins.

## 1. Introduction

The control of gene expression at the post-transcriptional level is critical for diverse biological processes including proper organismal development in multicellular organisms. Many regulators, including RNA-binding proteins (RBPs), act to control the stability of target mRNA transcripts through the recognition of key sequence elements in the 3′ UTRs of mRNAs [1, 2]. A recent survey of all known human RBPs indicated that a substantial fraction of human RBPs bind to mRNAs, however, for any given RBP, the binding specificity, set of mRNA targets, and functional role for the RBP at each target still remains poorly understood [3].

The PUF (Pumilio and FBF [fem-3 binding factor]) family of proteins represent one of the most well-studied classes of RBPs [1, 4, 5]. PUF proteins possess a shared C-terminal Pum homology domain (PUM-HD). Structurally, the human PUM-HD consists of 8 helical repeats containing specific amino acids that both intercalate and form hydrogen bonds and van der Waals contacts with target RNA, resulting in exquisite specificity for a UGUANAUA consensus sequence motif or PUM Recognition Element (PRE) [6, 7]. Recognition by the PUM-HD is modular and specificity for a given base can be changed through mutation of a set of three key amino acids in a single repeat [7, 8]. Furthermore, the sequence specificity by PUM-HD across species can be predicted from the identity of these three key amino acids across the helical repeats in any given PUM-HD [9]. Thus, there are slight differences in the exact set of sequences recognized by the PUM-HD of different PUF family members and, in addition, interactions with protein partners can alter sequence preference [10–12].

Functionally, the PUF family of proteins have been implicated in post-transcriptional regulation underlying control of developmental processes [1]. One of the founding members of the family, *Drosophila* Pum, together with the Nos protein, is needed for correct body patterning in the developing fly embryo [13, 14]. Patterning is accomplished by location-specific repression of the *hunchback* mRNA through sequence-specific recognition of a nanos response element (NRE) in the *hunchback* 3′ UTR [15]. In humans, there are two members of the PUF family, PUM1 and PUM2, which share 75% overall sequence identity with 91% sequence identity in the PUM-HD. In addition, human PUM1 and PUM2 share 78% and 79% sequence identity in the PUM-HD to DmPum, respectively [5, 16]. Human PUM1 and PUM2 are expressed across tissues and their expression is highly overlapping [5, 16] suggesting that they likely act redundantly. Mammalian PUM proteins have been implicated in spermatogenesis [17, 18], neuronal development and function[19–24], immune function [25, 26], and cancer [27–30]. PUM1 missense and deletion mutants lead to adult-onset ataxia (Pumilio1-related cerebellar ataxia, PRCA) and loss of one copy leads to developmental delay and seizures (Pumilio1-associated developmental disability, ataxia, and seizure; PADDAS) [31]. Yet, the targets responsible for these biological outcomes are largely opaque.

Targeted experiments have indicated that human PUM1 and PUM2 are capable of repressing expression of a luciferase reporter through recognition of PREs in the reporter gene’s 3′ UTR, likely through recruitment of the CCR4-NOT complex and subsequent degradation of the mRNA target [32]. Additionally, similar assays have shown that repression by the human PUM2 PUM-HD alone—that is lacking the N-terminal domains of PUM2—requires the polyA binding protein PABPC1, suggesting that the human PUMs could accelerate mRNA degradation by inhibiting translation [33]. However, PUM-mediated repression is not the only type of gene regulation by human Pumilio proteins. Recently, expression of a key regulator of hematopoietic stem cell differentiation, FOXP1, was shown to be enhanced by human PUM1/2 binding to the 3′ UTR [29]. Furthermore, measurements of changes in global steady-state RNA abundance between wild-type (WT) and PUM1/2 knockdown conditions have identified hundreds of RNAs that either increase or decrease in abundance upon PUM1/2 knockdown [34]. Follow-up experiments have confirmed activation of key targets by human PUMs through the use of a reporter gene-target 3′ UTR fusion construct [34], indicating that human PUMs directly activate some mRNA targets. However, the mechanism of PUM-mediated activation remains to be elucidated.

High-throughput measurements of PUM1 and PUM2 binding sites *in vivo* have confirmed high specificity for a PRE and have identified a diverse set of PUM targets in human cell lines, including those involved in regulating neuronal function and signaling cascades [35–38]. Thus, sequence-specific recognition of the PRE is an important aspect of target recognition for the PUM proteins. However, key questions about PUM-mediated gene regulation remain. There are on the order of 10,000 PRE sites across the full set of annotated human 3′ UTRs, but only ∼1000 genes change in steady state RNA levels under PUM1/2 knockdown [34]. Additionally, models using a simple count of PREs in the 3′ UTR of a transcript do not completely capture the complexity of PUM-mediated gene regulation [34]. The identification of additional sequence features that discriminate functional PREs from apparently non-functional PREs will improve the understanding of PUM-mediated gene regulation. Furthermore, as the measurement of steady-state RNA levels do not allow for differentiation between the individual contributions of transcription rates and RNA stability, we instead set out to directly measure changes in RNA stability under PUM1/2 knockdown conditions. Through the use of high-throughput sequencing methodologies, we demonstrate that human PUM1/2 modulate the abundance of mRNA targets primarily through controlling mRNA stability and not transcription rates. We demonstrate, through high-throughput *in vitro* binding assays, that PUM1 and PUM2 PUM-HDs have highly similar preferences for the same sets of sequences. Consistent with prior reports, we find that PUM1/2 control the mRNA stability of transcripts involved in signaling pathways, neuronal development, and transcriptional control. In addition, we identify a key set of contextual features around PREs that contribute meaningful information in predicting PUM-mediated regulation including proximity to the 3′ end of a transcript and the AU content around PRE sites. Taken together, our study illuminates key contributors to determining functional PRE sites and represents a rich resource for interrogating the control of mRNA stability by the PUM RBPs.

## 2. Results

### 2.1. Bru-seq and BruChase-seq reveal PUM-mediated effects on mRNA stability

In order to measure the effect of the human PUM1 and PUM2 proteins on mRNA stability at a transcriptome-wide scale, we employed the Bru-seq and BruChase-seq methodology [39]. In brief, Bru-seq and BruChase-seq involve the metabolic labeling of RNA using 5-bromouridine (BrU), which is readily taken up by the cells and incorporated into the nascent NTP pool [40]. After incubation with BrU over a short time period, newly synthesized and labeled RNAs are selectively pulled out of isolated total RNA using an anti-BrdU antibody and sequenced. Labeled RNA abundance is then tracked over time by continuing to grow the cells in the absence of BrU and isolating BrU-labeled RNA at additional time points. To distinguish relative changes in transcription rates from relative changes in RNA stability between WT and PUM1/2 knockdown cells, we chose two time points: (1) a zero hour time point taken at the transition to unlabeled media after 30 minutes of incubation in BrU-containing media and (2) at six hours, a time point chosen to coincide with the average mRNA half-life in cultured mammalian cells [41–43]. To determine the impact of PUM1/2 on relative RNA abundances, the experiment was performed in the presence of a mix of siRNAs targeting both *PUM1* and *PUM2* mRNAs (siPUM) or in the presence of scrambled non-targeting control siRNAs (NTC), as previously established [32, 34](Figure 1A). Cells were treated with siRNAs for 48 hours before BrU labeling, identically to the method used in Bohn et al. [34], to allow for PUM depletion prior to labeling. Overall, four biological replicate samples were collected for each time point and RNAi condition resulting in a total of 16 samples and above the minimum recommendations for replicates suggested by the ENCODE consortium for RNA-seq and ChIP-seq experiments [44, 45]. HEK293 cells were chosen for this study as they express both PUM1 and PUM2, have been previously used to analyze PUM activity [32, 34], support efficient BrU-labeling [46], and support RNA interference [47]. As we have previously demonstrated [32, 34], knockdown of both PUM1 and PUM2 is necessary to alleviate PUM repression of PRE-containing mRNAs. It is important to note that the use of two time points does not allow for determination of full decay rate constants for each transcript, but it does allow for measurements of relative changes in mRNA stability between the two conditions [48].

**Figure 1:**
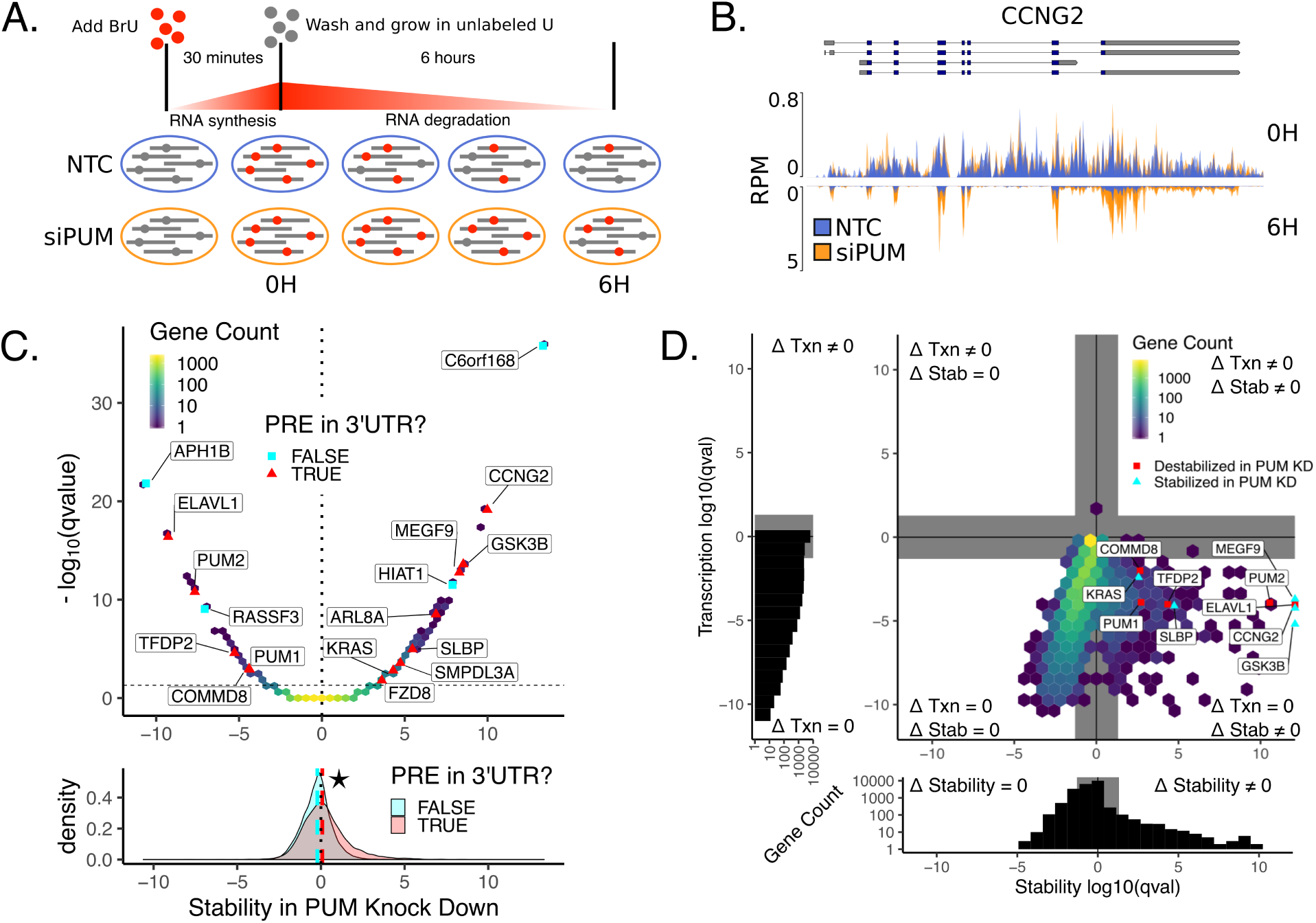
Bru-seq and BruChase-seq allow for determination of PUM-mediated effects on RNA stability. A) Experimental design for measuring PUM-mediated effects on RNA stability. HEK293 cells incubated for 30 minutes in the presence of 2mM BrU prior to time 0. Cells were then washed and cultured in media containing 20 mM unlabeled uridine for six hours. At 0 and 6 hour timepoints, a portion of cells were harvested and BrU labeled RNA was isolated for sequencing. Changes in relative RNA abundance between the 0 and 6 hour time points were compared between cells grown in the presence of silencing RNA targeting *PUM1* and *PUM2* (siPUM) and a non-targeting control siRNA (NTC). Cells were treated with siRNAs for 48 hours prior to BrU labeling to allow for PUM depletion. B) Read coverage traces for CCNG2 as measured in reads per million (RPM). Traces are shown for siPUM (orange) and NTC (blue) conditions at both 0H (top) and 6H (inverted bottom) time points. Four replicates for each combination of siRNA and time point are overlaid. Known isoforms for *CCNG2* are represented above. C) (Top) Volcano hexbin plot displaying global changes in RNA stability under PUM knockdown conditions. Stability in PUM knockdown is represented by a normalized interaction term between time and condition, where positive values indicate stabilization upon PUM knockdown and negative values indicate destabilization upon PUM knockdown (see Methods for details). No change in stability is represented with a dotted line at 0. Statistical significance at an FDR corrected p-value < 0.05 is represented with a horizontal dashed line. A selection of genes known to be regulated by PUM [34, 35] and genes newly identified in this study are labeled. For selected genes only, red triangles indicate genes that have a PRE in any annotated 3′ UTR as determined by a match to the PUM1 motif we identified using SEQRS (Figure 2A). Gray squares indicate genes that did not have a PRE in their 3′ UTR. Unlabeled genes are binned into a two-dimensional histogram to avoid overplotting. (Bottom) Marginal distribution of Stability in PUM knockdown for genes with a PRE in their 3′ UTR (red) and genes without a PRE in their 3′ UTR (gray). Median values for each distribution are plotted as a dashed line in the appropriate color. The star indicates a statistically significant difference in the median stability as measured by a two-sided permutation of shuffled labels (*n* =1000, p < 0.001). D) Analysis of changes in transcription vs. changes in stability. Four separate statistical tests were calculated for each gene: 1. a test for statistically significant changes in RNA stability (Δ Stability ≠ 0), 2. a test for statistically significant changes in transcription (Δ Txn ≠ 0), 3. a test for no change in RNA stability (Δ Stability = 0), and 4. a test for no change in transcription (Δ Txn = 0). Genes are plotted as an (x,y)-coordinate where each coordinate represents the ± log_10_(FDR corrected p-value) of the test with greater evidence (Δ ≠ 0, +*log*_10_; or Δ = 0, -log_10_) for each axis (see Methods for details). Representative genes displaying a range of stability effects are labeled. Red squares represent genes that were destabilized in PUM knockdown, whereas red triangles represent genes that were stabilized in PUM knockdown. All other genes were binned into a two dimensional histogram. Gray rectangles represented a statistical significance cutoff of q-value > 0.05. (Left and Below) Marginal histograms for each axis are plotted with matching gray rectangles to represent the same statistical significance cutoff of q-value > 0.05.

Clear changes in RNA abundance can be seen between time points and conditions at the gene level. Consider the Cyclin G2 (CCNG2) mRNA which encodes a cyclin involved in the cell cycle, contains 2 PREs in its 3′ UTR, and was among the most dramatically affected mRNAs (Figure 1B). At the 0 hr time point, read coverage resulting from recent transcription for four distinct replicates in each condition can be seen (Read coverage includes immature RNAs that still contain introns) (Figure 1B top). At the six hour time point, only mature RNA remains, with read coverage primarily observed at exons and no longer prevalent in the intronic regions (Figure 1B bottom). Here, silencing of both PUM1 and PUM2 clearly increases RNA abundance relative to the nontargeting control at the 6 hr time point, but does not appear to impact transcription as seen at the 0 hr time point.

To quantify the effect of silencing PUM1 and PUM2 on changes in relative labeled RNA abundance between the 0 and 6 hour time points, we used DEseq2 [49] to model the count of reads observed from each gene using a generalized linear model that considers the effects of time, condition, and the interaction between time and condition (see Methods for details). We interpret the term associated with the interaction between condition and time to be the PUM-mediated effect on stability—where a positive value indicates that an RNA was stabilized in the PUM knockdown condition and a negative value indicates that an RNA was de-stabilized in the PUM knockdown condition. Likewise, we interpret the condition term as the PUM-mediated effect on transcription rates, thus, we are able to separate the impacts of transcription from RNA stability using our experimental procedure and statistical methodology. We find that hundreds of genes show altered RNA stability under PUM knockdown conditions. Figure 1C displays an overview of PUM-mediated effects on stability as a volcano plot, with 12,165 genes represented in a two-dimensional histogram. Using an FDR-corrected p-value threshold of 0.05 and a fold-change cutoff of *log*_2_(1.75) (see Methods), we found 44 genes were statistically significantly de-stabilized (56 with no fold-change cutoff) and 200 genes were statistically significantly stabilized in the PUM knockdown condition (252 with no fold-change cutoff). Of these genes, 30 were also identified as having lower abundance under PUM knockdown in the Bohn et al. [34] RNA-seq data set (37 with no fold-change cutoff). Likewise, 95 were also identified as having higher abundance under PUM knockdown in the Bohn et al. [34] RNA-seq data set (106 with no fold-change cutoff). As expected, in our data both *PUM1* and *PUM2* were substantially destabilized in the PUM knockdown condition relative to the WT condition indicating that the siRNAs were successful in disrupting PUM1/2 expression and that our methodology is capable of detecting known changes in RNA stability. Additionally, we found that genes with a PRE in their 3′ UTR were, on average, more stabilized in the PUM knockdown condition than those without a PRE in their 3′ UTR (Figure 1C bottom). Taken together, this suggests that PUM1/2 are selectively modulating the RNA stability of target transcripts.

To further examine the effects of PUM knockdown on both transcription and stability, we tested for statistically significant changes under a null model centered around a log_2_ fold change of 0 for both the condition term (transcription) and the interaction between condition and time (stability). In addition, for each term, we also tested for a statistically significant lack of change by considering a null model centered around the boundary of a defined region of practical equivalence spanning from − log_2_(1.75) to log_2_(1.75)(see Methods for details); such a test is important because failure to reject the null hypothesis cannot, by itself, be taken as evidence favoring the alternative. In total, four statistical tests were run for each gene: a test for change and a test for no change for both transcription and stability. For each axis, the smaller of the two FDR-corrected p-values (i.e. test for change vs. test for no change) was chosen as the coordinate for that term, which enabled classification of each gene into one of four quadrants: 1. Genes that change in both stability and transcription (Figure 1D, upper right quadrant), 2. genes that change only in stability (Figure 1D, lower right quadrant), 3. genes that change only in transcription (Figure 1D, upper left quadrant) and 4. genes that change in neither (Figure 1D, lower left quadrant). Thus, using this methodology, we identified 213 genes with a statistically significant change in stability (Figure 1D lower right quadrant). We were also able to identify a set of 2,834 genes with evidence for no change in stability under our experimental conditions (Figure 1D lower left quadrant) and 19,744 genes we were have insufficient information to reliably classify. Additionally, we show only one gene, *ETV1*, with a statistically significant change in transcription, 11,527 genes with statistically significant lack of change in transcription and 11,263 genes we have insufficient information to reliably classify. Taken together and consistent with the Pumilio proteins’ role in post-transcriptional regulation, these results suggest that PUMs regulate gene expression at the level of RNA stability and not transcriptional initiation. Furthermore, this analysis allows us to divide the genes into those in which Pumilio knockdown has an effect on RNA stability and those in which there is evidence for a lack of effect on RNA stability, a stronger statement than simply failing to reject the null hypothesis that no change was occurring. The words EFFECT and NOEFFECT will be used to refer to these respective gene classes throughout the rest of the paper.

### 2.2. SEQRS shows conserved preference for the canonical UGUANAUA PRE by Pumilio proteins

The sequence preferences for both the full length PUM1 and PUM2 have been previously probed *in vivo* [36–38, 50] and the sequence preferences for the RNA-binding domains of both PUM1 and PUM2 were probed *in vitro* [10, 51, 52]. Each of these approaches and methodologies agree on a general preference for the UGUANAUA consensus motif for both PUM1 and PUM2, with subtle differences in the information content for the Position Weight Matrices (PWM)s obtained from each technique, particularly at the 3′ end of the PWM. However, prior *in vitro* determination of human PUM sequence preferences have involved only one round of selection [51] or a selected subset of possible sequences [52]. Thus, to compare the binding specificity of the PUM-HD of the human PUM1 and human PUM2 proteins we applied *in vitro* selection and high-throughput sequencing of <u>R</u>NA and <u>s</u>equence specificity landscapes (SEQRS) to purified PUM-HDs of each protein [53]. Similar to <u>s</u>ystematic <u>e</u>volution of ligands by <u>ex</u>ponential enrichment (SELEX) [54], SEQRS allows for the determination of an RNA-binding protein’s sequence specificity by selecting for RNAs that interact with the RBP out of a pool of random 20mers generated by T7 transcription of a synthesized DNA library. The RNA pulled-down from a previous round is reverse-transcribed into DNA to be used as the input for the next round of transcription and selection, allowing for exponential enrichment of preferred sequences for any RBP of interest. We applied five rounds of SEQRS to the PUM1 and PUM2 PUM-HDs separately and quantified the abundance for each of the 65536 possible 8mers in the sequencing libraries for each round (including 8mers that would overlap with the adjacent static adapter sequences see Methods for details).

To obtain representative PWMs for each round of selection (Figure 2A,B (top)), we used the top enriched 8mer, UGUAAAUA, as a seed sequence to create a multinomial model from the abundance of every possible single mismatched 8mer to the seed sequence (see Methods for details). This data analysis approach has yielded similar results to that of expectation-maximization algorithms such as MEME [55] and has been used successfully with SELEX experiments using DNA-binding proteins [56, 57]. We also applied this same analysis pipeline to previously published SEQRS analysis of the *D. Melanogaster* Pumilio PUM-HD [53] and find that it readily captures the *D. mel* Pum sequence preference for the canonical UGUANAUA PRE (Figure 2D (top)). However, the PWMs defined here (Figure 2A,B,D (top panels)) are representative of only the most highly enriched sequences in each dataset and round.

**Figure 2:**
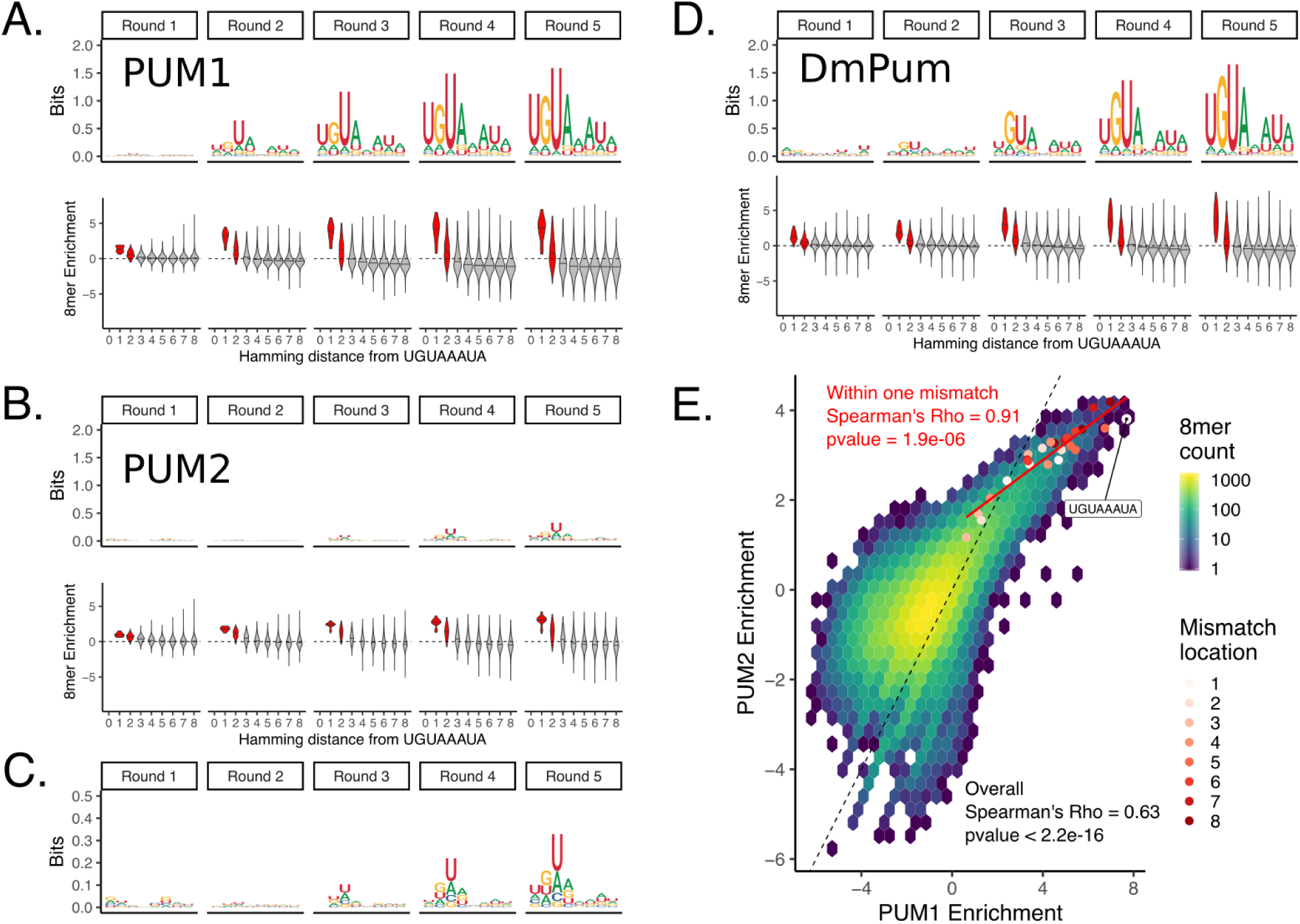
SEQRS analysis of Human PUM1 and PUM2 PUM-HDs reveals preference for the canonical PUM Recognition Element. A) (Top) Position weight matrices representing 8mer sequence preferences for purified Human PUM1 PUM-HD, as determined for each SEQRS round. (Bottom) 8mer enrichment, as measured by log_2_(Enrichment SE-QRS round/ Enrichment no protein) (see Methods for details) for each 8mer as binned by Hamming distance from the canonical UGUAAAUA PUM recognition element. Enrichment scores for 8mers within 2 mismatches are filled in red. B) Same as in A, but for Human PUM2 PUM-HD. C) Closer view of Human PUM2 PUM-HD PWMs. D) Same as in A, but for Drosophila Pum PUM-HD. E) Correlation of 8mer enrichment between Human PUM1 and Human PUM2 PUM-HDs. Enrichment for all possible 8mers are displayed in a two dimensional histogram. The dashed black line represents one to one correspondence. All 8mers within one mismatch to the UGUAAAUA sequence are plotted as red points with the color specifying the position within the motif where the mismatch occurs. The red line is a linear fit using only the UGUAAAUA 8mer and all 8mers within one mismatch.

In order to determine how representative the UGUANAUA consensus motif is for the entire dataset of each protein, we grouped each 8mer based on its similarity to the UGUAAAUA seed sequence as measured by the number of mismatches to that seed (Hamming distance). We then considered the relative enrichment of a given 8mer within each round compared to its relative enrichment within the input pool. Thus, scores above 0 indicate higher relative abundance than the input pool for a given 8mer and scores below 0 indicate lower relative abundance. Here, we see that 8mers within 1-2 mismatches of the UGUAAAUA seed sequence are highly enriched compared to 8mers with more than 2 mismatches across each round for each protein (Figure 2A,B,D (bottom)). However, the high level of variation in enrichment scores with higher numbers of mismatches and the inclusion of some 8mers with high enrichment scores in these groups, suggests that only considering sequences that are within 1 or 2 mismatches of the canonical PRE (here represented by UGUAAAUA) may not fully describe PUM binding specificity. Additionally, the PWM we obtained from our SEQRS experiment for PUM2 PUM-HD (Figure 2B-C) suggests that the PUM2 has much weaker enrichment for the canonical PUM PRE compared to PUM1, which is inconsistent with PUM2 sequence preferences obtained from *in vivo* transcriptome-wide experiments [36, 37]. This may indicate differences between *in vitro* and *in vivo* conditions that specifically impact PUM2 or may indicate that PUM2 PUM-HD does not bind as efficiently to RNA as the full-length PUM2 protein. However, comparing PWMs between these two proteins only considers the most highly enriched sequences in each dataset. As seen in Figure 2C, the consensus motif emerging from the PUM2 SEQRS data strongly resembles those for other PUMs, albeit with less apparent stringency. To compare the overall sequence preferences between PUM1 and PUM2 we plotted the enrichment scores for all possible 8mers in each dataset against each other (Figure 2E). We find that the 8mer enrichment scores between these two proteins are highly correlated (Spearman’s *ρ* = 0.63) which indicates that PUM1 and PUM2 PUM-HDs have overall similar sequence preferences when considering all possible sequences rather than highly enriched sequences. We also see that the PUM1 PUM-HD has an overall stronger enrichment for highly enriched sequences compared to PUM2, which may explain the differences in obtained PWMs for each protein. When considering only the 8mers within one mismatch to the UGUAAAUA seed sequence used for creating the PWMs, we find that enrichment scores between PUM1 and PUM2 are nearly perfectly correlated (Spearman’s *ρ* = 0.91). Furthermore, mismatches in the 3′ end of the motif appear to be less detrimental to enrichment by PUM1 and PUM2 compared to mismatches in the 5′ end of the motif, which is also represented by the lower information content at the 3′ end of the PWMs. Due to the overall similarity in sequence preferences between these two proteins and the higher overall information content in the PUM1 PWM, the SEQRS round 5 PWM for PUM1 will be used to determine PREs throughout the text, unless otherwise indicated.

### 2.3. Contextual features around PREs are associated with PUM-mediated RNA stability effects

Determining what distinguishes a functional binding site from a non-functional binding site is a major question for any RBP. Taken as a whole, RBPs tend to bind similar low sequence complexity motifs *in vitro* [51]. Additionally, probing of RBP binding *in vivo* at a transcriptome-wide scale, has indicated that the majority of predicted binding sites are not bound for some RBPs [58]. Global *in vivo* experiments with the Pumilio-family of proteins have established that mammalian Pumilio proteins recognize the UGUANAUA PRE in the 3′ UTR of target genes [22, 32, 36, 37]. However, predicting the PUM-mediated effect on gene expression from sequence information and/or PUM-binding measurements remains an elusive goal [34].

To determine sequence motifs *de novo* that have explanatory power for our RNA stability dataset, we used FIRE [59] to find motifs in the 3′ UTR of transcripts that share high mutual information with our RNA stability dataset by taking the normalized interaction term (see Methods for details) and discretizing it into ten bins, with an equal number of genes in each bin. Figure 3A shows that FIRE rediscovers the canonical UGUANAUA PRE using only the RNA stability data as input. Furthermore, the UGUANAUA PRE is enriched in transcripts that are highly stabilized under PUM knockdown conditions, suggesting that these transcripts are regulated by PUM through recognition of a UGUANAUA PRE in their 3′ UTR.

**Figure 3:**
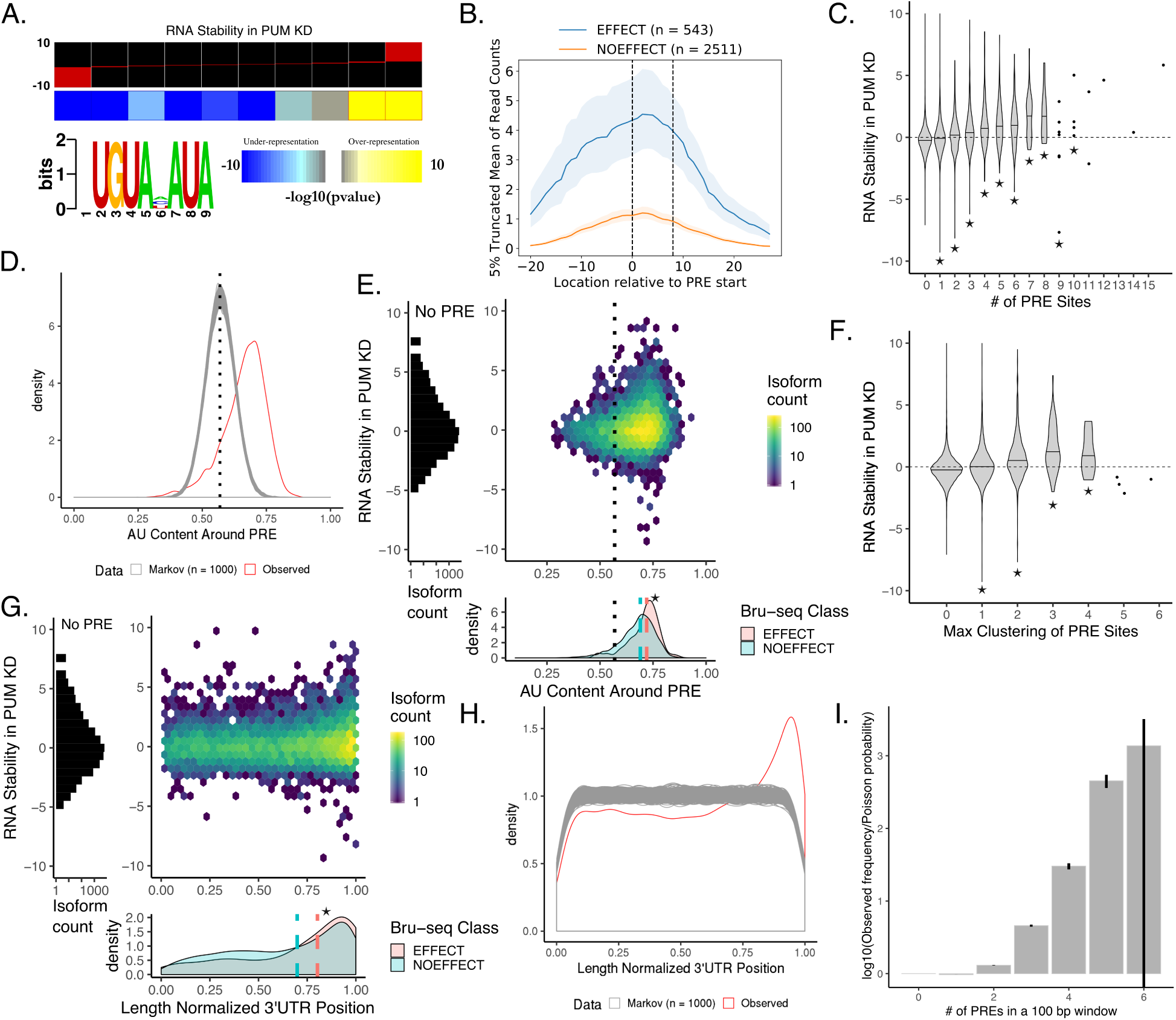
Features associated with a PUM Recognition Element (PRE) explain some variability in PUM-mediated effect on decay. A) Results of motif inference using FIRE [59] on the stability in PUM knockdown data discretized into 10 equally populated bins. Red bars within each bin represent the spread of RNA stability values within each bin. Stability in PUM knockdown is represented by a normalized interaction term between time and condition throughout this figure, where positive values indicate stabilization upon PUM knockdown and negative values indicate destabilization upon PUM knockdown (see Methods for details). B) 5% truncated average of Pum2 PAR-CLIP read coverage [37] over each PRE site in the 3′ UTRs of genes with a statistically significant change in RNA stability (blue) compared to genes in which there was a statistically significant lack of change in stability (orange; see Methods for details on NOEFFECT test). Shaded regions represent bootstrapping (n = 1,000) within each group. Dashed lines indicate the PRE site. C) Violin plots representing the distributions of RNA stability for genes with 0 to 15 PRE sites within their 3′ UTR. Stars represent statistical significance as measured by a Wilcoxon rank sum test using equality of pseudomedian with the 0 PRE case as the null hypothesis. D) Distribution of AU content in a 100 bp window around all unique PRE sites in the 3′ UTRs of the human transcriptome. The observed distribution (red) is compared to the distribution of AU content around PRE sites in 1,000 simulated sets of 3′ UTRs the same size as the true set of 3′ UTRs as simulated from a third order Markov model trained on the true 3′ UTR sequences. The dotted line represents the average overall AU content of the entire set of 3′ UTRs in the human transcriptome. E) Relationship of AU content in a 100 bp window around a PRE to RNA stability. (left) Marginal histogram of RNA stability for genes with 0 PREs in their 3′ UTRs. (right) 2D histogram of RNA stability and AU content around each PRE site for all genes with at least one PRE in the 3′ UTR. Dotted line represents the average AU content over the entire set of 3′ UTRs in the human transcriptome. (bottom) Marginal kernel density plot of AU content around a PRE site split amongst genes with a statistically significant change in RNA stability (red) and genes with a statistically significant lack of change in stability (blue). Dotted black line represents the average AU content of 3’UTRs. Dashed lines represent the median AU content around a PRE for the EFFECT (red) and NOEFFECT (blue) genes. The star represents a statistically significant difference in medians using a one-sided permutation test (n=1,000) of shuffled class labels. F) Violin plots representing the distributions of RNA stability for genes with 0 to 6 full PRE sites clustered within a 100 bp window. Stars represent statistical significance as measured by a Wilcoxon rank sum test using the 0 PRE case as the null distribution. G) Relationship of normalized location of PRE site in 3′ UTR to RNA stability. Plots as in (D). H) Distribution of length normalized locations of PRE sites in the 3′ UTRs of the human transcriptome. The observed distribution (red) is compared to that of PRE sites found in 1,000 simulated sets of 3′ UTRs calculated as in (G). I) Comparison of the observed frequencies of PRE site clustering over all possible 100 bp windows in the full set of human 3′ UTRs with at least 1 PRE in them to the probabilities expected from a Poisson null distribution. Error bars represent 95% confidence intervals based on 1,000 bootstraps of the observed distribution.

To determine whether there was evidence for PUM binding at PREs associated with a change in RNA stability, we used publicly available *in vivo* binding data for human PUM2 obtained using photoactivatable ribonucleoside-enhanced crosslinking and immunoprecipitation (PAR-CLIP) [37]. The PAR-CLIP technique involves incorporation of 4sU into the total cellular RNA pool allowing for efficient crosslinking of proteins that bind near an incorporated 4sU. Upon creation of sequencing libraries from PAR-CLIP samples, a T → C mutation is induced at the crosslinking site which can be used as additional evidence for a protein binding. We used PAR-CLIP data from Hafner et al. [37] to determine the amount of binding signal at PREs associated with transcripts that have a statistically significant change in RNA stability under PUM knockdown (EFFECT class, Figure 1D) and compared it to transcripts with a statistically significant lack of change in RNA stability (NOEFFECT class, Figure 1D). In Figure 3B, we report the average PAR-CLIP read coverage in a 40 bp window around PREs in the 3′ UTR of transcripts associated with the EFFECT and NOEFFECT classes. We use a 5% truncated mean to remove the impact of extreme outliers on the average coverage reported. To estimate a 95% confidence interval on the average coverage (shaded region), we performed bootstrapping (n = 1,000) by sampling vectors of read coverage for individual PREs with replacement. Here, we clearly see that PREs in transcripts with a change in RNA stability have higher binding signal than those with no change in RNA stability. This is consistent with higher overall PUM binding at PREs associated with changes in RNA stability but, as the PAR-CLIP signal is not normalized to RNA abundance, the possibility that these transcripts were simply more abundant under the PAR-CLIP conditions cannot be definitively ruled out.

We have shown that a PRE in the 3′ UTR is associated with a change in RNA stability under PUM knockdown and that PREs in transcripts with a change in RNA stability have evidence for being bound by PUM *in vivo*. However, knowledge of the presence or absence of a PRE in the 3′ UTR alone is not sufficient to predict the magnitude of PUM-mediated repression, and a wide variation in the effect of knocking down human PUM1 and PUM2 on steady-state RNA levels has been observed in previous transcriptome-wide analysis [34]. Here, we demonstrate that a similar level of variation can be seen in measurements of RNA stability. Figure 3C displays the overall distribution of RNA stability measurements for transcripts with increasing numbers of PREs in annotated 3′ UTRs. We find that an increase in the number of PREs is, on average, associated with an increase in RNA stability under PUM knockdown conditions compared to transcripts that do not have a PRE in their 3′ UTR. However, wide variations in RNA stability can be seen for each category, consistent with previous measurements of changes in steady state RNA levels under PUM knockdown [34]. Thus, a simple count of PREs does not fully explain PUM-mediated action at a particular transcript.

To explore the local sequence context around PREs, we trained a 3rd order Markov model on the full set of unique annotated human (hg19) 3′ UTRs that were greater than 3 basepairs long (29,380 3′ UTRs). Using this Markov model, we simulated 1,000 different sets of 29,380 3′ UTRs that were the same length and shared similar sequence composition to the set of true 3′ UTRs. We then searched for matching PREs in the simulated sets of 3′ UTRs and calculated the AU content in a 100 bp window around these PREs. On average, we discovered 12200 matching PREs (standard deviation of 112) in simulated sets of 3′ UTRs compared to the 14086 matching PREs in the annotated set of 3′ UTRs. We find that the true set of PREs have, on average, higher local AU content than PREs in simulated sets of 3′ UTRs (Figure 3D). Additionally, in the simulated 3′ UTRs the local AU content for PREs is centered around the average AU content for all 3′ UTRs, as would be expected if there was no selective pressure for PREs to occur in AU rich areas of 3′ UTRs. This analysis is consistent with Jiang et al. [60] who also observed a preference for PREs to occur in AU rich areas as compared to shuffled PREs with preserved overall sequence content. Here we further show that the local AU content surrounding a PRE is associated with a functional effect on PUM-mediated regulation.

To determine the relationship between local AU content and changes in RNA stability upon PUM knockdown, we plotted the AU content of a 100 bp window surrounding a PRE within a gene’s 3′ UTR against the corresponding RNA stability measurement for that gene (Figure 3E top). For 3′ UTRs with more than one PRE, the PRE with the highest local AU content was considered. We find that large changes in RNA stability are associated with higher local AU content. Additionally, PREs in transcripts that had a statistically significant stability effect in PUM knockdown had higher local AU content compared to PREs in transcripts with no change in stability (p < 0.001, Figure 3E bottom). These data indicate that local sequence context beyond the PRE plays a role in PUM function.

Previously proposed mechanisms of PUM-mediated control of RNA stability involve interaction with the CCR4-NOT complex and/or PABPs, both of which act at the 3′ end of mRNA transcripts to promote deadenylation or participate in translation initiation [32, 33]. Thus, the location of PUM binding sites within the 3′ UTR of target transcripts may play a role in determining PUM-mediated effects on stability by physically locating PUM near known co-regulators. Using the Markov models described above, we also determined the location of PREs within 3′ UTRs. As shown in Figure 3H, we observe that the observed distribution of true PRE locations in length-normalized 3′ UTRs appear enriched towards the 3′ end of 3′ UTRs (red) as compared to PREs found within 1000 simulated sets of 3′ UTRs (gray). Again, this suggests a selective pressure for PRE sites to exist at the 3′ end of 3′ UTRs as compared to the uniform distribution of PREs found in simulated 3′ UTRs with similar sequence properties. Like the AU content analysis, this analysis is also consistent with observations made by Jiang et al. [60] who saw an enrichment towards the 3′ end for PRE locations in the full set of human 3′ UTRs compared to a shuffled PRE motif with preserved overall sequence content. While these approaches are complementary, our approach allows for the exact identity of the PRE to remain intact thereby maintaining a PRE-centric assessment rather than one based solely on the general sequence content within the motif. Additionally, we observe that transcripts with a PRE towards the 3′ end of the 3′ UTR tend to have a larger RNA stability effect (Figure 3G center) and PREs in transcripts that had a statistically significant change in stability in PUM knockdown were, on average, closer to the 3′ end of the 3′ UTR than those with no change in RNA stability (p < 0.001, Figure 3G bottom), suggesting a functional role for PRE location in the 3′ UTR of target transcripts.

High throughput analysis of many human RBPs has indicated that some RBPs prefer to bind bipartite motifs, suggesting that clustering of RBP binding sites may contribute to binding specificity and subsequent function [51]. To determine the relationship between PRE clustering and RNA stability in PUM knockdown, we discretized transcripts according to the maximum number of complete PREs that were within a sliding 100 bp window in the 3′ UTR of a transcript and plotted the distribution of RNA stability measurements for each cluster (Figure 3F). Similar to the association with the number of PREs (Figure 3C), we find that having more PREs clustered together is associated, on average, with a higher stabilization effect under PUM knockdown conditions. We also find that PREs tend to cluster together more than one would expect by chance by determining the divergence from a simple Poisson model (Figure 3I, p < 0.001 for clusters 2-5; see Methods for details). Taken together, this analysis suggests that clustering of PREs may facilitate PUM action on target transcripts.

### 2.4. Pumilio proteins modulate the stability of genes involved in neural development, cell signaling, and gene regulation

Mammalian Pumilio proteins have been shown to regulate a diverse set of genes, including those involved in signaling pathways, transcriptional regulation, and neurological functions [18, 22, 23, 34, 35]. Consistent with prior observations, we see changes in RNA stability for genes involved in these functions. For example, multiple epidermal growth factor-like-domains 9 (*MEGF9*) is a transmembrane protein that is highly expressed in the central and peripheral nervous system and its expression appears to be regulated during nervous system development in mice [61]. We see strong stabilization of the *MEGF9* transcript under PUM knockdown conditions (Figure 4A top). Furthermore, of the five PREs we identify in two unique 3′ UTRs for *MEGF9*, we see the most PUM2 binding signal for the 3′-most PRE (Figure 4A bottom right). Additionally, we see that the 3′-most PRE has high local AU content compared to the overall distribution of PRE sites (Figure 4A bottom left). Taken together, these data implicate the PUM proteins as direct post-transcriptional regulators of *MEGF9*.

**Figure 4:**
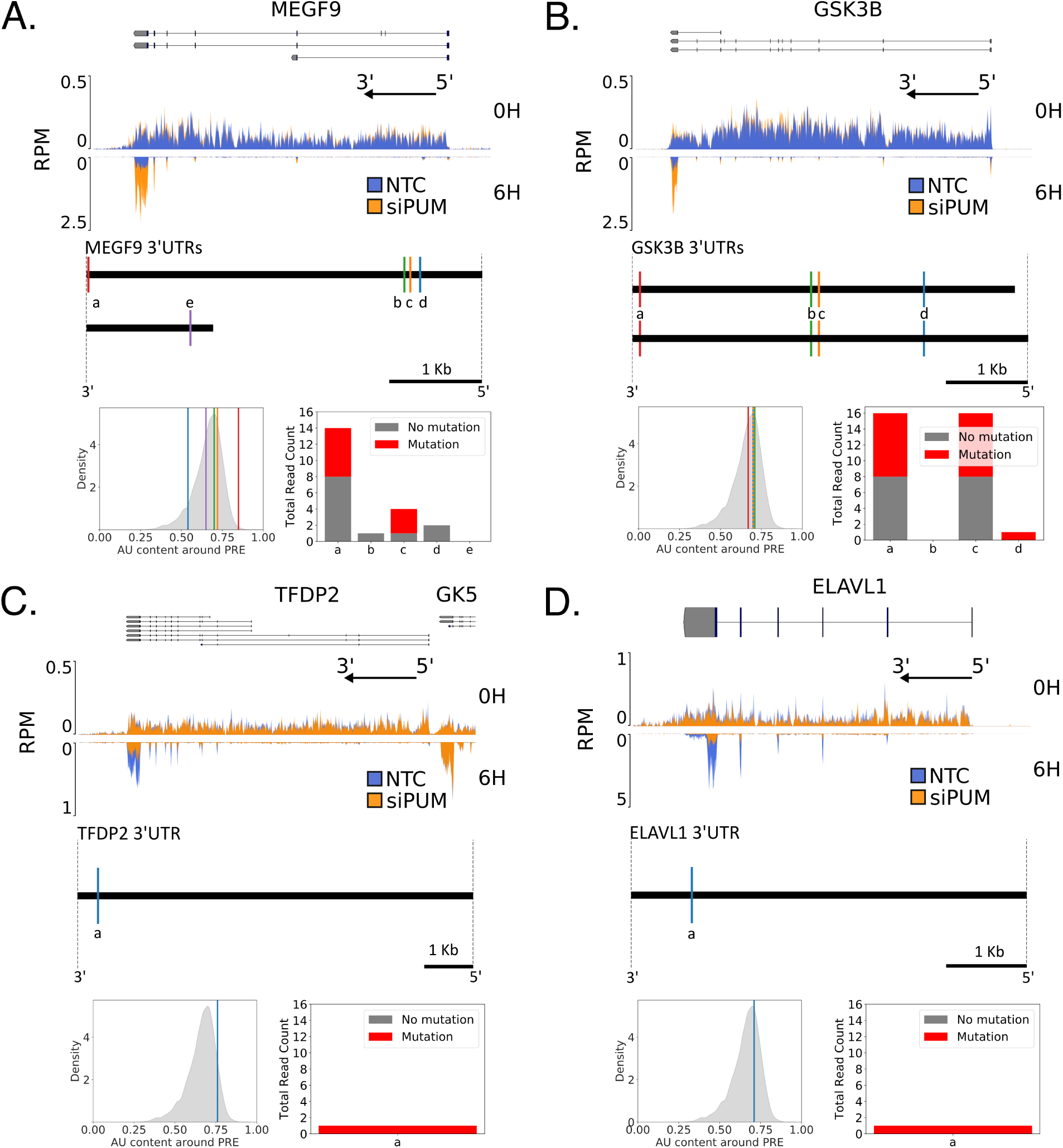
PUM-mediated effects on RNA stability under PUM knockdown include stabilization and destabilization. A) (top) Read coverage traces for *MEGF9* and surrounding region (chr9:123348195-123491765, hg19) as measured in reads per million (RPM). Traces are shown for siPUM (orange) and NTC (blue) conditions at both 0H (upper track) and 6H (inverted lower track) time points. Four replicates for each combination of siRNA and time point are overlaid. Known isoforms for *MEGF9* are represented above. The black arrow indicates the direction of the 5′ and 3′ ends of the transcribed RNA molecule from the gene shown. (Below) Diagram of unique *MEGF9* 3′ UTRs. Sites matching the PUM1 SEQRS motif are represented as vertical lines and labeled alphabetically from 3′ to 5′ for each UTR. (Below left) AU content of a 100 bp window around each PRE labeled above in the overall distribution of surrounding AU content for all PUM1 SEQRS motif matches in the entire set of 3′ UTRs. (Below right) PAR-CLIP read coverage [37] of 40 bp around each indicated PRE. Number of reads with a T→C mutation are shown in red, whereas the number reads with no T→C mutation are shown in gray. B) As in A), but for *GSK3B* and surrounding region (chr3:119509500-119848000). C) As in A), but for *TFDP2* and surrounding region (chr3:141630000-141900000). Annotations for the 3′ end of the *GK5* gene are included due to their proximity to the *TFDP2* 5′ end. D) As in A), but for *ELAVL1* and surrounding region (chr19:8015000-8080000).

Another transcript that is strongly stabilized under PUM knockdown conditions is glycogen synthase kinase-3 B (*GSK3B*) (Figure 4B top). GSK3B is a serine-threonine kinase that is involved in the regulation of diverse cellular processes and its misregulation is associated with neurological disease [62, 63]. We identify four PREs in *GSK3B* 3′ UTRs (Figure 4B below) with largely similar adjacent AU content (Figure 4B bottom left). We also find that the 3′ most distal PRE has evidence for PUM2 binding consistent with the global trends we describe in Figure 3. Like *MEGF9*, this evidence suggests that PUM proteins are involved in destabilizing *GSK3B* transcripts.

We also see examples of RNAs that are destabilized when PUM is knocked down, suggesting that PUM may actually act to stabilize these transcripts under conditions containing WT levels of PUM expression. Transcription dimerization partner 2 (*TFDP2*) encodes a protein that cooperates with E2F transcription factors to regulate genes important for cell cycle progression; dysregulation of this system can lead to cancer [64]. PUM proteins have been previously shown to regulate another member of the E2F family by functionally cooperating to enhance the effect of miRNA-mediated regulation of *E2F3* expression [65]. Furthermore, regulation of *TFDP2* by the liver-specific miRNA *miR-122* has been shown to be important for preventing up-regulation of *c-Myc* in hepatic cells [66]. We observe that *TFDP2* is highly destabilized under PUM knockdown conditions (Figure 4C top). Additionally, we find that the *TFDP2* 3′ UTR has a single PRE site toward the 3′ end of the 3′ UTR and has high adjacent AU content (Figure 4C bottom and lower left). However, there is limited evidence for PUM2 binding in PAR-CLIP data (Figure 4C lower right). One possible mechanism for PUM mediated activation of *TFDP2* is by acting to block regulation by miRNAs; however, the nearest conserved miRNA site of a conserved miRNA family to the PRE is over 100 bases away [67] and further evidence would be needed to establish this link.

Another example of a highly destabilized transcript under PUM knockdown conditions is the embryonic lethal abnormal vision 1 (*ELAVL1*) or HuR RNA-binding protein (Figure 4D top). The *ELAVL1* RBP stabilizes RNA transcripts by binding to AU-rich elements in the 3′ UTR of transcripts [68] and its dysregulation is associated with several different types of cancer [69]. We found one PRE in the 3′ UTR of *ELAVL1* (Figure 4D bottom). This motif is found towards the 3′ end of the 3′ UTR but has average local AU enrichment compared to other PREs found across all annotated 3′ UTRs (Figure 4D lower left). Additionally, there is limited evidence for binding by PUM2 at either of the PREs in the *ELAVL1* 3′ UTR (Figure 4D lower right). Taken together, this suggests that *ELAVL1* may be indirectly regulated by PUM.

To discover categories of genes that are globally associated with RNA stability changes in PUM knockdown, we applied iPAGE—a computational tool that uses mutual information to find informative Gene Ontology (GO) terms associated with discretized gene expression data [70]—to our stability dataset as represented by the normalized interaction term discretized into 5 equally populated bins. It is worth noting that this analysis will discover pathways regulated both indirectly and directly by PUM out of the full set of annotated GO terms. Figure 5A displays the iPAGE results with several GO terms that are either significantly overrepresented (red-filled box) or underrepresented (blue-filled box) across the full range of stability data. We see several enriched GO term categories that are consistent with previous reports of changes in steady-state RNA levels under PUM knockdown in HEK293 cells [34] including categories related to guanyl-nucleotide exchange factor activity (GO:0005085), WNT signaling (GO:0030177), nucleosome (GO:0000786) and platelet-derived growth factor receptor signaling (GO:00048008).

**Figure 5:**
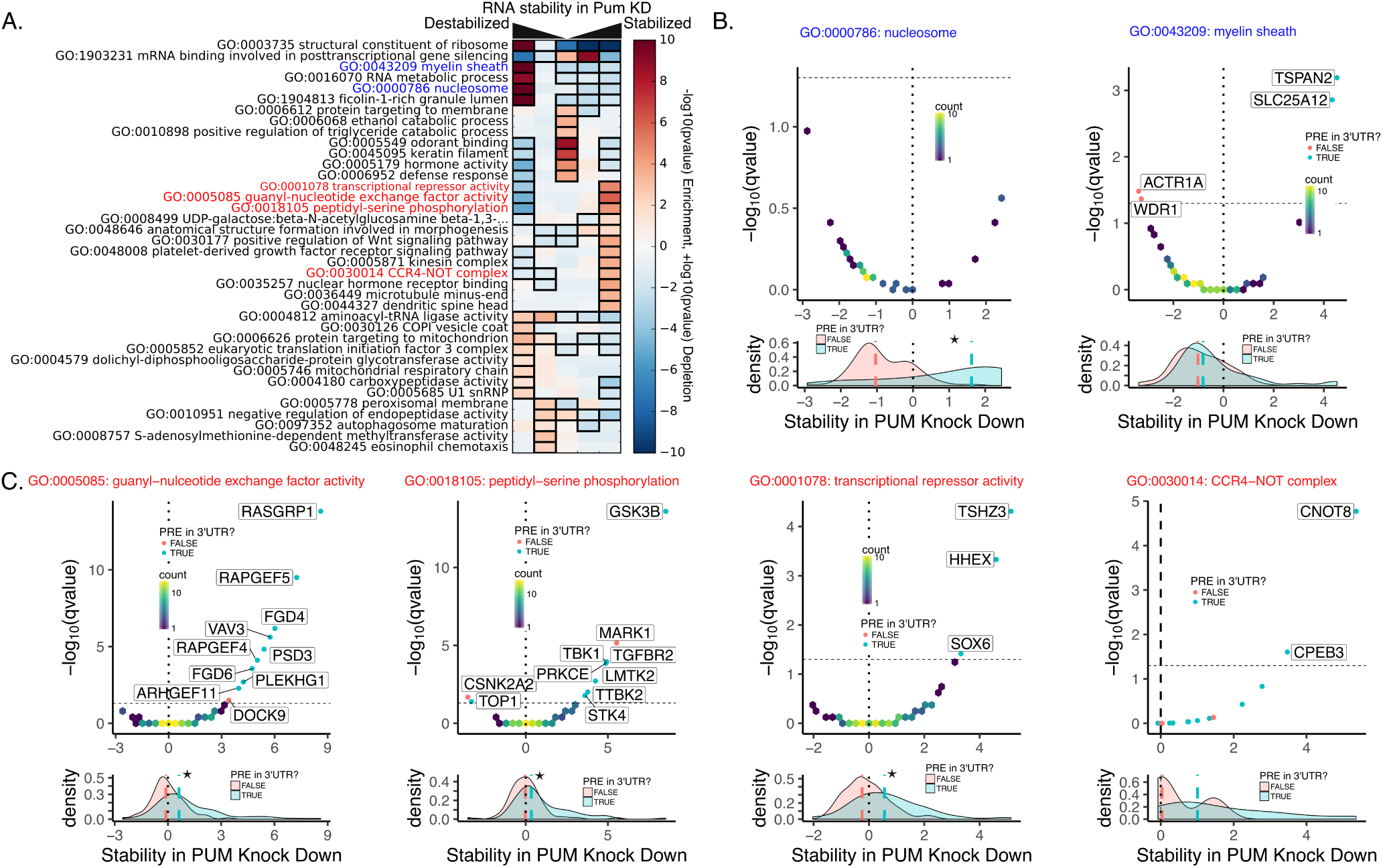
Gene ontology terms associated with PUM-mediated changes in RNA stability. A) Results of iPAGE analysis to find GO terms sharing mutual information with RNA stability discretized into 5 equally populated bins. Red bins indicate over representation of genes associated with the corresponding GO term. Blue bins indicate under representation of genes associated with the corresponding GO term. A black box indicates a statistically significant over or under representation with a p-value < 0.05 using a hypergeometric test [70]. Throughout this figure stability in PUM knockdown is represented by a normalized interaction term between time and condition, where positive values indicate stabilization upon PUM knockdown and negative values indicate destabilization upon PUM knockdown (see Methods for details). B) Selected GO terms whose members are over represented in the RNAs that are destabilized under PUM knockdown, as labeled in blue in panel A. For each GO term, a volcano plot is shown for all genes within the GO term. Volcano plots are shown as two dimensional histograms for genes below a statistical significance threshold (q-value < 0.05) and as individual points for genes above the statistical significance threshold. Individual points are blue if a PRE can be found within any annotated 3′ UTR for that gene and red otherwise. The dashed line represents the statistical significance threshold and the dotted line represents no change in RNA stability under PUM knockdown. Below each volcano plot is a marginal density plot for the RNA stability split into two categories within the specified GO term: Genes with a PRE in any annotated 3′ UTR (blue) and genes with no PRE in any annotated 3′ UTR (red). Medians for each distribution are shown as dashed lines in the appropriate color. The black dotted line represents no change in RNA stability, as in the volcano plot above. A star represents a statistically significant (p < 0.05) difference in the medians as tested by a two-sided permutation test of shuffled group labels (n = 1000). C) As in (B), but for selected GO terms whose members are over represented in the RNAs that are stabilized under PUM knockdown, as labeled in red in panel A.

For a finer grain view, we plotted the RNA stability results for each gene involved in selected GO terms as indicated by either blue (destabilized in PUM KD) or red (stabilized in PUM KD) text for that GO term in Figure 5A. In Figure 5B, we show two selected GO terms whose members tend to be de-stabilized upon PUM knockdown: nucleosome (GO:0000786, left) and myelin sheath (GO:0043209, right). For genes related to the nucleosome, we see a general destabilization under PUM knockdown conditions. However, when comparing genes within this GO term that have a PRE in their 3′ UTR to those that do not, we see that genes with a PRE in their 3′ UTR have a median stability upon PUM KD that is significantly higher than those without a PRE in their 3′ UTR (p < 0.001), suggesting that the destabilization of most nucleosome genes under PUM knockdown conditions may be mediated indirectly. Some of these effects could be explained by perturbation of the stem-loop binding protein (SLBP), as SLBP is a protein involved in the proper maturation of replication-dependent histone mRNAs [71], and we observe that *SLBP* is significantly stabilized under PUM knockdown conditions (Figure 1C).

PUM knockdown also causes a general de-stabilization of genes categorized into the myelin sheath GO term. A role for PUM in controlling the stability, either indirectly or directly, of genes involved in the myelin sheath is consistent with the previously identified role of mammalian PUMs in neurogenesis and neurodegenerative diseases [19, 22, 23, 31]. However, we see no evidence for a difference in stability between genes that have a PRE in their 3′ UTR compared to genes that do not have a PRE in their 3′ UTR. Furthermore, the genes that have a statistically significant de-stabilization under PUM knockdown have no PRE in their 3′ UTR, whereas the genes with a significant stabilization do, suggesting a complex role of PUM in modulating the stability of genes in this GO term, possibly arising mainly through indirect effects.

In Figure 5C, we report specific GO terms that were enriched in genes that were stabilized under PUM knockdown and thus likely contain many classic, PUM-repressed targets. Consistent with this idea, we find that each of these GO terms represent classes of genes that have previously been associated with PUM-mediated regulation. For instance, the guanyl-nucleotide exchange factor activity GO term (GO:0005085; Figure 5C, far-left) includes guanine nucleotide exchange factors (GEFs) which activate Rho-family GTPases to regulate a diverse suite of cellular functions, including cell-cycle progression, the actin cytoskeleton, and transcription [72]. Additionally, genes involved in peptidyl-serine phosphorylation (GO:0018105; Figure 5C, mid-left), represent a broad class of kinases, including those involved in neurological disease and inflammation [63, 73]. Finally genes involved in transcriptional repressor activity (GO:0001078, Figure 5C, mid-right), include proteins involved in regulating hematopoiesis and controlling neurological development [74–76]. Supporting the idea that PUMs are directly repressing subsets of genes within these GO terms we find that, for each GO term above, genes with a PRE in their 3′ UTR are significantly more stabilized under PUM knockdown than those with no PRE.

Of particular interest is the mild enrichment of genes that were stabilized under PUM knockdown for the CCR4-NOT complex GO term (GO:0030014; Figure 5C, far-right). Almost every gene in this GO term was stabilized under PUM knockdown to some extent. Although the overall effect of a PRE for genes in this category did not meet our threshold for statistical significance, several of the genes have a PRE in their 3′ UTR including both genes with a statistically significant change in stability. Human Pumilio proteins have been shown to interact with the CCR4-NOT complex and recruit the complex to target mRNAs for de-adenylation [32]. These data suggest that PUM could also be acting to directly inhibit CCR4-NOT expression and thus globally lower deadenylation rates, perhaps providing a feedback loop that further regulates PUM activity.

Overall, we observe that genes associated with GO terms that are stabilized under PUM knockdown have a significant association with PREs suggesting that these GO terms contain mainly genes that are direct targets of PUM. In contrast, we find that genes associated with GO terms that are destabilized under PUM knockdown do not have a significant association with PREs, suggesting that these GO terms contain mainly genes that are indirect targets.

### 2.5. Conditional random forest models allow for prediction of PUM-mediated effects from sequence-specific features

A long standing goal in the study of RBPs is to predict that RBPs effect on a given transcript from known features about possible targets. Previous models of PUM-mediated regulation have reported modest performance based on the number of PREs in various locations across the transcript including the 5′ UTR, CDS, and 3′ UTR [34]. Here, we use a different approach, which allows us to include a larger feature set of possible predictors for PUM-mediated regulation. Using conditional random forest models [77], we divided genes into EFFECT and NOEFFECT classes, as shown in Figure 1D. We used four different definitions for a PRE, (Figure 6A) including the SEQRS motifs we defined for PUM1 and PUM2 in Figure 2A-B, the PUM2 motif determined from Hafner et al. [37], and a regular expression (regex) representing UGUANAUW as defined from the PUM consensus sequence which has been used extensively to define PREs in previous publications [7, 34, 60]. We focused our analysis on PREs found in the 3′ UTRs of target genes. For each definition of a PRE, we calculated several features based on our analysis in Figure 3, including AU content around a PRE, clustering of PREs, total count of PREs, a score for PRE match to the specific PRE definition, relative location of the PRE in the 3′ UTR, number of miRNA sites near a PRE, and predicted secondary structure around a PRE. In addition to these features, we included motif matches for additional human RBPs, *in vivo* PUM binding data, predictions of secondary structure, and the fraction optimal codons for the CDS of target genes (see Methods for details). As our data is highly unbalanced (199 EFFECT genes and 2535 NOEFFECT genes, after only including genes that are present in all features) we trained 10 different machine learning models where the NOEFFECT class was randomly downsampled to match the number of EFFECT class genes in each model. Within each downsampled dataset, 5-fold cross validation was performed to assess performance.

**Figure 6:**
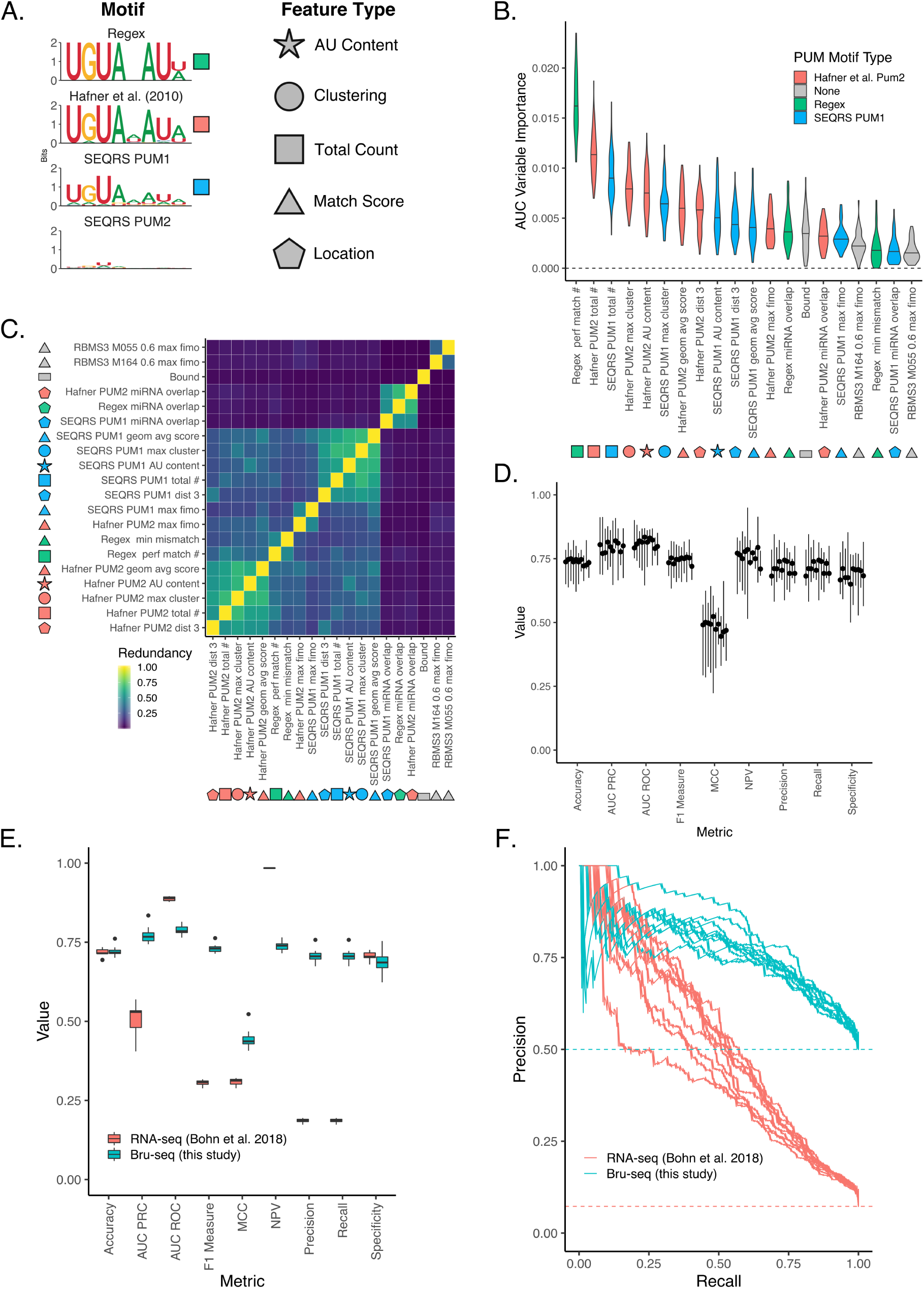
Predicting PUM-mediated effect on decay using both sequence-based and experimental features. A) Motifs used to calculate features for machine learning. Shapes indicate the type of feature calculated, whereas colors indicate the motif used to calculate those features. Total count is a simple count of motifs; Match score refers to a numerical value indicating how well a sequence matches a motif; clustering indicates motif proximity to additional instances of the same motif; location indicates features associated with a single motif’s location on the 3′ UTR. Shapes filled in with the appropriate color are used to label features throughout the rest of the figure. B) Variable importance plot displaying the top twenty most important features, as determined by training a conditional random forest classifier on PUM decay data (see Methods for details including information on feature names). Violin plots represent density from ten separate downsamplings of the majority class, each with five fold cross-validation. An AUC based variable importance measure is used as described in Janitza et al. [78]. C) Calculation of the redundancy in information between the top twenty most important variables, as determined in A. Redundancy is calculated in the information-theoretic sense (see Methods for details) where 1 is completely redundant information and 0 is no redundancy in information between the two variables. D) Cross-validation of conditional random forest classifier performance. Each boxplot represents a separate downsample of the majority, no PUM-mediated effect class. Values for each boxplot represent the performance metric as calculated for each of five folds using a classification cutoff of 0.5. E) Performance of conditional random forest models on the steady state RNA data-set from [34]. Blue boxplots represent values from seperate downsamplings of the majority, no PUM-mediated effect class used to train the model on the Bru-seq and BruChase-seq data set. Red boxplots indicate values from testing each model on the Bohn et al. steady-state RNA-seq data set. Metrics were calculated using a classification cutoff of 0.5. F) Precision Recall curves using the models in E. Each line represents one of ten conditional random forest models trained on separate down sampled sets of the entire Bru-seq and BruChase-seq data set and tested on the steady state RNA data set.

To determine which features best help predict EFFECT genes from NOEFFECT genes, we used an AUC-based permutation variable importance measure [78], which indicates the average change in the area under the curve (AUC) of a receiver operator characteristic (ROC) plot across all trees with observations from both classes in the forest when the predictor of interest is permuted. By permuting the feature of interest and measuring the change in AUC of the ROC curve, one can measure the importance of that variable in predictive performance. Typically values of the AUC of a ROC curve span from 0.5 to 1.0 where 1.0 indicates perfect classification performance and 0.5 indicates random guessing of class distinctions. Since the AUC-based variable importance measure is calculated using the change in AUC when the predictor is permuted, the expected values are much smaller and fall between 0.0 and 0.06 in simulated cases with 65 predictors and variable numbers of observations from n=100 to n=1,000 [78]. Higher values indicate a larger drop in performance when that variable is permuted; thus, the variables can be ranked based on their unique contribution to the model, with higher values indicating a more important individual contribution. Figure 6B displays the top 20 variables ranked according to their average AUC-based variable performance across all 50 models (10 sets of downsampled models with 5-fold cross-validation each). Count based metrics enumerating the total number of PREs within the 3′ UTR appear to be the most important variable for predicting a PUM-mediated effect in the Bru-seq and BruChase-seq data. In addition, local AU content and PRE clustering appear to be substantial contributors to the models. To a lesser extent, the number of miRNA sites around a PRE, the location of the PRE in the 3′ UTR, and the “Bound” status of the 3′ UTR also appear to contribute meaningfully to our models. It is possible that each of these variables contain largely the same information (i.e., whether or not the 3′ UTR has a PRE or not in it). Thus, in order to rule out the possibility that each feature was simply differentiating between genes with a PRE in their 3′ UTR from genes without a PRE, we trained separate models for each motif definition where we only considered genes that have at least one PRE present in their 3′ UTR. Each of these models also displayed substantial contributions for AU content, clustering, and total count in predicting PUM-mediated regulation, as measured by Bru-seq and BruChase-seq (Figure S2A-D left panel) suggesting that each of these features are contributing meaningful information to the model.

The high similarity in appearance between each of the definitions of a PRE we include here led us to explore how much redundant information is contained between each of the top 20 highest contributing features. To measure redundancy, we use an information theoretic definition based on discretization of each feature (see Methods for details). In Figure 6C, we display the redundancy between the top 20 features as a hierarchically clustered heatmap, where a value of 1.0 indicates that the features contain exactly the same information and a value of 0.0 indicates that the features share no information. Here, we can see that features that are defined around the same motif definition or feature-type tend to share information (as expected). However, despite their similarity in appearance, there are some differences in information content between the different motif definitions and different feature types, indicating that there is knowledge to be gained outside of a simple PRE count.

To assess the performance of our conditional random forest models we considered several typical performance measures including summary metrics (Accuracy, F1 measure, Matthews correlation coefficient [MCC], Area Under the Curve of a Precision-Recall Curve [AUC PRC], and AUC ROC), and metrics more focused on performance for positive or negative cases (Negative Predictive Value [NPV], Precision, Recall, Specificity). We considered each of these metrics for all 50 models (10 downsampled datasets with 5-fold cross-validation each) at a classification probability cutoff of 0.5. The full range of values obtained are displayed in Figure 6D. It is evident that the models are robust to both downsampling and cross validation and the performance hovers around 0.75 for each metric (and 0.5 for MCC), indicating balanced performance in predicting both positive and negative classes. These results are robust even in the case where we only use one PRE definition and only consider genes that contain a PRE in their 3′ UTR (Figure S2A-D).

In order to determine the predictive efficacy of our models we tested their performance against the Bohn et al. [34] RNA-seq dataset which was not used to the train the models (Figure 6E). Here, the performance on the trained Bru-seq and BruChase-seq data is reported as the five-fold crossvalidation performance for each of the 10 downsampled models. To observe the overall performance of the models, we display precision-recall curves on both the Bru-seq and BruChase-seq data on which the model was trained and the RNA-seq data for each of the 10 different models (Figure 6F). The baseline is defined separately for each dataset as the overall class balance between the positive and negative class. A perfect model tends toward the upper right of the graph, and a poor model follows the dotted baseline for that dataset. Despite the differences in technique and biological implications between RNA-seq and Bru-seq and BruChase-seq in determining PUM-mediated gene regulation, we find that the models trained on Bru-seq and BruChase-seq are able to perform well in predicting PUM-mediated regulation in RNA-seq data. We see similar performance when considering a single definition for a PRE and only considering genes that have a least one PRE in their 3′ UTR (Figure S2A-D). Although the features we have included here are not sufficient to fully describe PUM-mediated gene regulation in human cells, we have demonstrated a clear functional association and predictive utility for PUM motifs (i.e. match scores and count of PREs) as well as contextual features around PREs including the location, neighboring AU content, clustering of PREs, and overlap with predicted miRNA sites.

## 3. Discussion

Through the combination of our high-throughput probing of RNA decay and the mining of sequence information in the 3′ UTRs of human transcripts, we were able to establish several general rules of PUM-mediated gene regulation in human cells.

### 3.1. Human PUM proteins control gene expression by modulating RNA stability

Previous studies have established that both PUM1 and PUM2 control the stability of individual transcripts through recognition of a UGUANAUA PRE [32]. Transcriptome-wide measurements in PUM1 and PUM2 knockdown conditions have shown that hundreds of RNAs change in abundance, as measured using RNA-seq [34]. However, measurements of RNA abundance using RNA-seq only allow for determination of changes in steady-state RNA abundances and do not allow one to differentiate effects from changes in RNA stability versus changes in transcription rates. Through the use of metabolic labeling, we are able to differentiate the effects of knocking down both PUM1 and PUM2 on transcription from the effects on RNA stability [39]. Our results indicate that perturbing the expression of human PUM1 and PUM2 has a widespread effect on the mRNA stability of many transcripts in HEK293 cells, but does not appear to perturb transcription rates in any meaningful way, as measured by our system. Rather than determine full decay rate constants for each transcript, which would have required the use of additional time points throughout the chase period of our experiment, we chose to determine relative changes in RNA stability using just two time points. The measurements obtained from these experiments cannot be interpreted on an absolute scale, but the rank order of stability measurements within the experiment is preserved, allowing us to determine the relative effects of PUM knockdown between any two genes [48]. Consistent with the changes in steady-state RNA levels determined under PUM knockdown conditions, we see transcripts that are both destabilized and stabilized. As expected, the number of genes that are stabilized under PUM knockdown is much higher than the number of genes that were destabilized, which is consistent with PUM’s role in reducing the expression levels of target genes likely through the recruitment of the CCR4-NOT complex and subsequent destabilization of the transcript [32].

### 3.2. General rules for predicting PUM-mediated activation remain elusive

In contrast with the clear and robust effects of PUM on PUM-repressed transcripts, the mechanism for the rarer case of PUM-mediated stabilization remains unclear. Measurements using luminescent reporter assays have shown activation of a subset of predicted PUM-activated transcripts that is dependent on the presence of a PRE in the 3′ UTR of the reporter [34]. Furthermore, direct binding of PUM1 or PUM2 to PREs present in the *FOXP1* 3′ UTR has been reported to promote expression of the FOXP1 protein, an important regulator of the cell cycle in hematopoietic stem cells [29]. Conversely, when considering PAR-CLIP measurements of PUM2 occupancy at PREs for only the transcripts that were destabilized under PUM knockdown, we find inconclusive evidence for binding in targeted examples (Figure 4C,D) and an insufficient number of examples to draw firm conclusions when considering the group as a whole separately from the stabilized transcripts (data not shown). Furthermore, attempts to classify transcripts that were stabilized in PUM knockdown from those that were destabilized using random forest models with identical feature sets to those used in Figure 6 showed poor performance, possibly due to the small number of examples for transcripts that were destabilized under PUM knockdown. There is also the possibility that the destabilization of the transcripts under PUM knockdown are indirect effects mediated through another factor that PUM is either directly regulating or PUM is competing with for binding. It is likely that the PUM-mediated activation of genes found through high-throughput studies represent a combination of direct and indirect targets. However, despite the clear evidence for direct PUM-mediated activation of some targets, general rules for predicting PUM-mediated activation remain elusive and mechanistic insights into PUM-mediated activation of key targets will require further study.

### 3.3. PUM1 and PUM2 have shared sequence preferences

Using SEQRS [53] on purified PUM-HDs for both PUM1 and PUM2, we find a strong preference for the UGUANAUA motif for PUM1 and, somewhat surprisingly, a much weaker preference for this motif for PUM2. However, when considering the enrichment of all possible 8mers, we see that the preferences for each PUM-HD are highly correlated with a larger magnitude in enrichment for PUM1 PUM-HD compared to PUM2 PUM-HD. Our approach uses a random library of RNA sequences to determine RNA binding preferences and our analysis of PUM1 qualitatively agrees with previous *in vitro* approaches with randomized libraries [51]. However, using a curated library of sequences based on mutations from the consensus UGUANAUA motif, Jarmoskaite et al. [52] created a thermodynamic model for PUM2 binding that considers the effects of non-consecutive bases in target recognition, as opposed to our simpler model that only considers the frequency of occurrence of consecutive bases in a fully randomized library. Using this model, they show that PUM-HDs from both PUM1 and PUM2 share nearly identical sequence preferences, which is in agreement with our strong correlation in enrichment between the two proteins.

When we considered the local sequence content and location of PREs, we found that PREs tend to be located towards the 3′ end of the 3′ UTR and have high local AU content. We are not the first to observe these properties, as Jiang et al. [60] also arrived to this conclusion by comparing the locations of shuffled PREs. However, we instead considered the locations of PREs in simulated sets of 3′ UTRs that share similar trinucleotide content to that of the true set of 3′ UTRs and this strengthens the claim that PREs are enriched in these areas more than one would expect by chance. Furthermore, we are able to connect these observations directly to functional outputs, showing that PREs in transcripts that had a significant change in RNA stability under PUM knockdown are closer to the 3′ end of the 3′ UTR and have higher flanking AU content, suggesting a functional role for the location of PREs within the 3′ UTR itself. The non-random propensity of PREs to occur towards the 3′ end of the 3′ UTR is consistent with a model where PUMs recruit the CCR4-NOT complex for de-adenylation of target sequences.

### 3.4. Human Pumilio proteins regulate genes involved in signaling pathways

When looking at the classes of genes that are stabilized under PUM knockdown, we find that many GO terms with evidence for direct repression by PUMs revolve around regulating signaling pathways mediated by proteins including kinases (GO:0018105), GEFs (GO:0005085), and receptor signaling (GO:0030177, GO:0048008). The role of mammalian Pumilio proteins in modulating signaling through controlling mRNA levels has been well established. In human testes, PUM2 is thought to interact with DAZL proteins to regulate germ-line development and many GTP-binding, receptor-associated, and GEF encoding-mRNAs are found among a list of targets that co-immunoprecipitate with both proteins [17]. Similarly, PUM1 has been shown to be important in mouse testis development through downregulation of many proteins involved in MAPK signaling and ultimate activation of p53 [18]. In fact, it has been argued that an ancestral function of the PUF family of proteins is to regulate the maintenance of stem cells and cells that behave in a stem cell-like manner through the down-regulation of kinases involved in critical signaling pathways [1]. Many studies looking at mRNAs associated with PUM1 or PUM2 binding in mammalian cells tend to find similar sets of GO terms overlapping with PUM bound targets. Early RIP-Chip experiments with human PUM1 and PUM2 found that genes bound by both proteins belonged to GO terms associated with the Ras pathway, MAPK kinase cascade, PDGF signaling pathway, WNT signaling pathway, small GTPase-mediated signal transduction, and transcription factor activity, among others [35, 36]. More recent iCLIP experiments in mouse brains have found that mouse PUM1 and PUM2 bind transcripts for genes associated with WNT signaling, regulation of MAP kinase activity, small GTPase-mediated signal transduction, and several categories related to neural development [22]. Similarly, changes in steady-state RNA abundance under both human PUM1 and human PUM2 knockdown identified several similar classes of genes including WNT signaling, GEF activity, NOTCH signaling, and PDGF signaling [34]. Each of the categories noted above is consistent with identified biological roles for mammalian PUMs. For example, mice lacking PUM1 and PUM2 have impaired learning and memory, as well as decreased neural stem cell proliferation and survival [22]. Further, human PUM1 haploinsufficiency is associated with developmental delay and ataxia [31]. Likewise, PUM2-deficient mice are more prone to chemically-induced seizures and have impaired nesting abilities [20], and mouse PUM2 regulates neuronal specification in cortical neurogenesis [23]. Our work shows that genes in these GO categories are modulated at the level of mRNA stability, likely through direct interaction of the human PUM proteins by recognition of PREs in the 3′ UTR of transcripts.

In many ways, post-transcriptional regulation of proteins involved in signaling cascades is an ideal way to rapidly modulate those pathways. In contrast to the delay in time between the control of mRNA synthesis and the resulting protein production involved in regulating a gene at the transcriptional level, post-transcriptional regulation allows for a rapid dampening of expression levels directly where protein synthesis is occurring. Furthermore, gene regulation in the cytosol allows for the possibility of localized control of expression [79]. In fact, temporal and localized control of gene expression—important for proper development of the fly embryo—was exactly how the PUF family of proteins were initially discovered [13]. Given the emerging role for human PUM proteins in neuronal development and function, and the need for localized control of gene expression in neuronal tissue [80] it is conceivable that PUM proteins could be heavily involved in RNA polarity within the neuron as has been observed in *C. elegans* olfactory neurons [81].

### 3.5. Prediction of PUM-mediated regulation defines a set of general principles for an ideal PUM target site

Many attempts have been made to predict gene regulation by Pumilio proteins given sequence information about the possible targets. Previously, a biologically inspired model based strictly on the count of PREs within the 5′ UTR, CDS, and 3′ UTR was fit to steady state RNA levels [34]. In this model, the effects of having multiple PREs on a single transcript were found to be less than linear on the target response to PUM knockdown, which was interpreted to indicate that multiple PRE sites function to increase the odds of having a PUM bound and that a single PRE likely performs most of the functions needed for PUM-mediated regulation [34]. In this study we expanded the feature set of possible predictors for PUM-mediated activity and determine a set of rules that define a functional PRE. Consistent with the Bohn et al. [34], we find that a simple count of PREs in the 3′ UTR acts as the best predictor for PUM activity. However, surprisingly we find that the simple UGUA.AU[AU] regular expression outperforms more sophisticated PWM-based definitions from either *in vivo* and *in vitro* high throughput data. This may indicate that, although PUMs can bind PREs with mismatches from this consensus motif, the UGUANAUA may represent the “ideal” PRE for functional regulation. In fact, structural studies of human PUM1 and PUM2 have identified three different modes of binding between the nucleotide bases of the fifth base in the consensus motif and the amino acids of PUM repeats 4 and 5. Lu and Hall [82] show that changes between these modes of binding do not alter PUM binding affinity, but could conceivably present different surfaces for effector proteins. Although our regular expression allows for any base at the fifth position, PUM repeats are modular [7] and it is conceivable that a similar mechanism could apply to other bases in the motif. Additionally this suggests that PUM binding to the UGUANAUA consensus motif could represent the ideal structure for PUMs interaction with effector molecules. We also find sequence features surrounding a PRE to be important in predicting PUM activity on a target. High AU content and position within the 3′ UTR both appear to be important for predicting mammalian PUM regulation. Consistent with prior reports of cooperativity between PUM and miRNAs [36, 50, 60, 65], we find that a count of predicted miRNA sites near PREs helps predict PUM effect, with a higher number of miRNA sites near a PRE indicating a larger stabilization under PUM knockdown (Figure S1A). It is possible that PUM could act to block or enhance miRNA function through direct interactions with the miRNA machinery or through local rearrangements of RNA secondary structure.

Secondary structure has been predicted to have an effect on many RBPs [51] and PUM has been shown to change secondary structure upon binding to facilitate miRNA interaction [65]. However, we found that *in silico* predictions of RNA secondary structure around PREs were not predictive of PUM function (Figure S1C). Targeted regression models considering PRE count and structure performed worse when structural information was added (data not shown). Recent studies have shown that structural probing experiments used in tandem with *in silico* folding algorithms vastly improve biological predictions based on structural information [83]. Similar methods may be needed to determine the role of secondary structure in PUM-mediated regulation. Alternatively, PUM proteins may be able to overcome RNA secondary structure in order to bind PREs, in which case, secondary structure would have no bearing on PUM binding. Similarly, RNA modifications may limit the ability for PUM to recognize PREs. Recent efforts have identified m6A sites across the human transcriptome at single nucleotide resolution [84]; however, we find limited to no overlap between m6A sites and PREs (data not shown).

There has also been a recent interest in the role of codon optimality in mRNA decay in human cells [85, 86]. Using, as a measure of codon optimality, the fraction of optimal codons—where a codon is designated as optimal if its Codon Stability Coefficient is positive [87]—we find that PUM targets undergoing PUM-mediated decay in our data set have a lower fraction of optimal codons on average than those with no PUM-mediated effect (Figure S1B). However, the fraction of optimal codons did not rank in the top twenty most important features for differentiating between transcripts subject to PUM-mediated decay from those that are not affected in our machine learning models (Figure 6). Recent studies have implicated codon optimality as an important determinant of mRNA stability in eukaryotes [85–88] and it is conceivable that PUM proteins could be directly mediating some of these effects. However, it is also possible that RNAs with a lower fraction of optimal codons represent more ideal targets for PUM or that PUM could be interacting with the factors that mediate decay for RNAs with less optimal codons. Further studies will be needed to establish the relationship between PUM and codon optimality.

By combining high-throughput functional data with statistical modeling, we have identified several contextual features around PREs that have improved our understanding of PUM-mediated gene regulation and increased our ability to predict PUM targets. However, there is still substantial room for improvement. Recent successes in Pumilio target prediction in *Drosophila* have come from characterizing binding partners of DmPum: Nos and Brat [89]. Nos binds together with DmPum to modulate the 5′ sequence specificity of the Pum-Nos complex, thus introducing finetune control over Pum target recognition [11]. A recent study identified many new and previously known interacting partners for the human PUM1 and PUM2 proteins including DAZL, PABP, FMRP, miRISC, and members of the CCR4-NOT complex [90]. Like the Nos/DmPum example, these partners likely add an additional layer of information in the control of PUM-mediated gene regulation. Furthermore, the probing of RNA secondary structure *in vivo* may allow for better incorporation of secondary structural information into models of PUM-mediated regulation. Finally, we were unable to find determinants of PUM-mediated activation, an area that is rich for future targeted experiments.

## 4. Materials and Methods

### 4.1. Experimental methodology

#### 4.1.1. SEQRS protein purification

Methods are reproduced here from Weidmann et al. [11]. Recombinant Halo-tag PUM1 RBD (aa 828-1176) and Halo-tag PUM2 RBD (aa 705-1050) were expressed from plasmid pFN18A (Promega) in KRX *E. coli* cells (Promega) in 2xYT media with 25 *µ*g/mL kanamycin and 2mM MgSO_4_ at 37°C to OD_600_ of 0.7–0.9, at which point protein expression was induced with 0.1% (w/v) rhamnose for 3hr. The PUM RBD expression constructs were originally described in Van Etten et al. [32]. Cell pellets were washed with 50mM Tris-HCl, pH 8.0, 10% (w/v) sucrose and pelleted again. Pellets were suspended in 25mL of 50mM Tris-HCl pH 8.0, 0.5mM EDTA, 2mM MgCl_2_, 150mM NaCl, 1mM DTT, 0.05% (v/v) Igepal CA-630, 1mM PMSF, 10 *µ*g/ml aprotinin, 10 *µ*g/ml pepstatin, and 10 *µ*g/ml leupeptin. To lyse cells, lysozyme was added to a final concentration of 0.5 mg/mL and cells were incubated at 4°C for 30min with gentle rocking. MgCl_2_ was increased to 7mM and DNase I (Roche) was added to 10 *µ*g/mL, followed by incubation for 20 min. Lysates were cleared at 50,000 × *g* for 30min at 4°C. Halo-tag containing proteins were purified using Magnetic HaloLink Resin (Promega) at 4°C. Beads were washed 3 times with 50mM Tris-HCl pH 8.0, 0.5mM EDTA, 2mM MgCl2, 1M NaCl, 1mM DTT, 0.5% [v/v] Igepal CA-630) and 3 times with Elution Buffer (50mM Tris-HCl, pH 7.6, 150mM NaCl, 1mM DTT, 20% [v/v] glycerol).

To confirm protein expression, beads were resuspended in Elution Buffer with 30 U of AcTEV protease (Invitrogen), cleavage proceeded for 24hr at 4°C, and beads were removed by centrifugation through a micro-spin column (Bio-Rad). Concentration of eluted protein was measured by Bradford assay, followed by coomassie stained SDS-PAGE analysis.

SEQRS was conducted on PUM1 PUM-HD and PUM2 PUM-HD as described in Campbell et al. [9] with minor modifications including the use of Magnetic Halolink beads (Promega). The PUM test proteins remained covalently bound via N-terminal Halotag to the beads.

The initial RNA library was transcribed from 1*µ*g of input dsDNA using the AmpliScribe T7-Flash Transcription Kit (Epicentre). 200 ng of DNase treated RNA library was added to 100 nM of Halo-tagged proteins immobilized onto magnetic resin (Promega). The volume of each binding reaction was 100*µ*l in SEQRS buffer containing 200 ng yeast tRNA competitor and 0.1 units of RNase inhibitor (Promega). The samples were incubated for 30min at 22°C prior to magnetic capture of the protein-RNA complex. The binding reaction was aspirated and the beads were washed four times with 200*µ*l of ice cold SEQRS buffer. After the final wash step, resin was suspended in elution buffer (1mM Tris pH 8.0) containing 10 pmol of the reverse transcription primer. Samples were heated to 65°C for 10min and then cooled on ice. A 5*µ*l aliquot of the sample was added to a 10*µ*l ImProm-II reverse transcription reaction (Promega). The ssDNA product was used as a template for 25 cycles of PCR using a 50*µ*l GoTaq reaction (Promega).

#### 4.1.2. Bru-seq and BruChase-seq experimental procedure

Bru-seq and BruChase-seq were conducted as described in Paulsen et al. [39] in HEK293 cells grown in the presence of siPUM1/2 or siNTC. RNAi conditions and siRNA sequences were previously described by Bohn et al. [34] and include treatment with siRNAs for 48hrs to allow for PUM depletion prior for BrU labeling. Four replicates were gathered for each time point and siRNA condition, resulting in 16 total samples. Resulting cDNA libraries were sequenced using an Illumina HiSeq 2000 via the University of Michigan Sequencing core.

### 4.2. Bru-seq and BruChase-seq Computational analysis

#### 4.2.1. Modeling PUM-mediated RNA decay

Sequencing reads were aligned to the human genome (hg19) and processed according to Paulsen et al. [39] up to obtaining read counts for exons and introns for each gene and sample. Our experimental design resulted in four different replicates of siNTC (WT) and siPUM1/2 (PUMKD) conditions with two different time points each: *t*_0*hr*_ and *t*_6*hr*_. For the *t*_0*hr*_ time points, read counts from both exons and introns were pooled for each gene. For the *t*_6*hr*_ time points, only read counts from exons were used. Read abundance was modeled using DESeq2 [49]. As described in Love et al. [49], DESeq2 models read count abundance *K* for gene *i* in sample *j* using the generalized linear model described below:

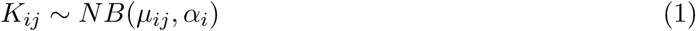

Where *α*_*i*_ is a gene-specific dispersion parameter for gene *i* and *µ*_*ij*_ is defined by the following:

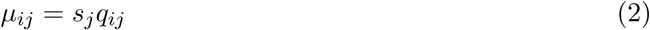

Here, *s*_*j*_ is a sample specific size factor used to put read count abundances on the same scale between samples. Finally, *q*_*i,j*_ is defined according to our design matrix:

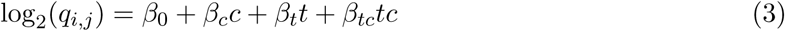

Where, *c* is an indicator variable that is 0 when the sample is in condition WT and 1 when the sample is in condition PUMKD. Likewise, *t* is an indicator variable that is 0 when sample is in the 0 hour time point and 1 when the sample is in the 6 hour time point. We interpret the *β*_*tc*_ term to represent changes in RNA stability resulting specifically from the PUM KD condition. Similarly we interpret the *β*_*c*_ term to represent changes in transcription rates between the two conditions. Throughout the text, unless otherwise noted, we report *β*_*tc*_ normalized by the reported standard error for the coefficient, which amounts to the Wald statistic computed for that term by DESeq2. Thus, the Wald statistic for the interaction term is denoted as “RNA stability in PUM KD” throughout the text and is a unitless quantity.

#### 4.2.2. Analysis of transcriptional vs. stability effects

To test for significant changes in transcription or stability, the Wald test statistic for the appropriate term—*β*_*c*_ for transcription and *β*_*tc*_ for stability—was calculated as described above. The Wald statistic was compared to a zero centered normal distribution and a two-tailed p-value was calculated using statistical programming language R’s pnorm function (n.b. this is virtually equivalent to the p-values calculated by the DEseq2 package for contrasts [49]). To test for a statistically significant lack of change in transcription or stability, the Wald statistic for the appropriate term was compared to a normal distribution centered at the nearest boundary of a region of practical equivalence (ROPE) and a two-tailed p-value was calculated using R’s pnorm function. The ROPE was defined as log_2_(1*/*1.75) – log_2_(1.75) and was chosen to be within the range of fold expression change of a RnLuc reporter gene with between one and three PREs in its minimal 3′ UTR [34]. Each p-value was FDR-corrected using the Benjamini-Hochberg procedure [91] and, for each term, the smaller of the two FDR-corrected p-values was reported. In order for a gene to be classified in the EFFECT class the following conditions had to be met: 1. its change in stability q-value had to be smaller than its no change in stability q-value; 2. Its change in stability q-value had to pass a cutoff of 0.05 for statistical significance; and 3. The original log_2_ fold-change value had to be outside the defined ROPE. In contrast, in order for a gene to be classified in the NOEFFECT class the following conditions had to be met: 1. it was not classified as an EFFECT gene; 2. its no change in stability q-value had to be smaller than its change in stability q-value; 3. its no change in stability q-value had to pass a cutoff of 0.05 for statistical significance; and 4. The original log_2_ fold-change value had to be within the defined ROPE. Genes not passing the criteria for either the EFFECT or NOEFFECT groups are those for which we lack sufficient information to make any strong statement on the effects of PUM knockdown.

### 4.3. SEQRS Computational analysis

Each raw sequencing read from the SEQRS experiments has the following expected structure: NNNNNN-CTGATCCTACCATCCGTGCT-NNNNNNNNNNNNNNNNNNNN-CACAGCTT CGTACCGAGCGG-GATCGGAAGA-XXXXXX-ATCTCGTA

Where X represents a known barcode sequence used to split the reads from a multiplexed experiment and N represents a random variable base. The *in vitro* transcription reaction uses the above sequence as a template resulting in RNA with sequence starting from the 3′ end of the CACAGCTTCGTACCGAGCGG downstream of the 20mer and going in the opposite direction. Thus, the RNA molecules in the SEQRS experiments are the reverse complement of the following: CTGATCCTACCATCCGTGCT-NNNNNNNNNNNNNNNNNNNN-CACAGCTTCGTACC GAGCGG

Raw sequencing reads were split by barcode, allowing for up to two pairwise mismatches on both the upstream and downstream adapter sequences. The 20mer variable regions and constant flanking adapter sequences of each read were reverse complemented and broken into all possible 8mer sequences using a sliding window, and raw counts for all possible 8mer abundances for each sequencing round for each protein were calculated using custom python scripts. For 8mers that overlapped the constant flanking adapter sequences, only 8mers that had at least one base in the variable region were considered.

To determine position-weight matrices that best represented selection by the protein of interest for that round, we followed the approach of Jolma et al. [57] in the analysis of DNA-binding proteins using SELEX. Briefly, a seed sequence is determined from the most abundant N-mer within that round. From this seed sequence, the abundance of each base at a given position was tallied when all other positions match the seed sequence. The PWM frequencies were determined by dividing each column of the resulting count matrix by its column sum. For all PWMs determined by this method we used a UGUAAAUA seed sequence. Unlike Jolma et al. [57] we do not include the correction for non-specific carryover of nucleic acid from the previous cycle as the assumption that no more than 25% of 8mers would be expected to be bound may not hold for RNA-binding proteins due to their promiscuous binding [51]. Instead, we accounted for the bias of the initial sequencing pool by calculating a PWM for the initial pool using the UGUAAAUA seed sequence. We then divided the position frequency matrix of each PWM by the initial sequencing pool’s position frequency matrix. Finally, we determined the bias-corrected frequency matrix by dividing each column of the matrix by its column sum.

In order to compare 8mer selection between rounds or proteins, the enrichment of a particular 8mer was calculated with the following equation:

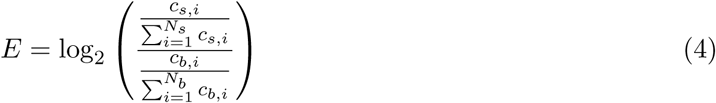

Where *c*_*s,i*_ represents the count for 8mer *i* in sample *s* and *c*_*b,i*_ represents the count for 8mer *i* in blank round where the input sequences were sampled. The DmPum data and corresponding blank sample was accessed from Weidmann et al. [11] and only the first five rounds were considered.

### 4.4. GO term analysis and iPAGE

GO term analysis was performed using the integrative pathway analysis of gene expression (iPAGE) software package [70]. Genes were discretized by the interaction term Wald test statistic into five-equally populated bins and iPAGE was run with default settings.

### 4.5. Determination of matching PREs

The full set of 3′ UTRs for hg19 genome was downloaded using the TxDb.Hsapiens.UCSC.-hg19.knownGene, BSgenome.Hsapiens.UCSC.hg19, and GenomicFeatures R packages. Matches to a given PWM across all 3′ UTRs were determined using the FIMO package with a uniform background using default cutoffs for reporting matches [92]. For PRE-centric figures, such as the heatmaps and violin plots in Figure 3 and Figure S1, each unique 3′ UTR isoform is matched to its corresponding “RNA stability in PUM KD” value by gene name, and each feature’s value is reported as the given summary statistic over a given 3′ UTR isoform for that feature, as described in the section below (i.e., for AU content, the value reported is the maximum AU content around any given PRE within that 3′ UTR isoform).

For *de novo* discovery of informative motifs in our Bru-seq and BruChase-seq dataset, we applied the finding informative regulatory elements (FIRE) software [59] with default settings to each unique 3′ UTR isoform matched to its “RNA stability in PUM KD” value and discretized into ten equally populated bins.

To calculate the location and AU content of PREs in randomly generated sets of the 3′ UTRs, a third order Markov model was trained on the annotated set of unique 3′ UTR isoforms from the hg19 genome. One thousand randomly simulated sets of 3′ UTRs—each with the same length as the annotated set of 3′ UTRs—was then generated using custom python scripts. For each of the thousand simulated sets of 3′ UTRs, the fifth round SEQRS PUM1 (Figure 2A) was used to search for matches using FIMO as described above. Here each individual PRE was considered in the calculation of the kernel density plots shown in Figure 3.

To determine the PAR-CLIP read coverage at identified PRE sites in the set of known unique 3′ UTR isoforms, raw reads were downloaded from SRA with accession numbers SRR048967 and SRR048968. Raw fastq files were processed with trimmomatic [93] and cutadapt [94] to remove low quality reads and illumina adapters. Processed reads were aligned to the hg19 genome using the STAR aligner with default parameters [95]. Read coverage and T to C mutations were determined for reads within 20 bp of each PRE in each unique 3′ UTR isoform for both EFFECT and NOEFFECT genes, individually, using custom python scripts. Coverage over all PREs was aligned and the bottom and top 5% of read coverage at each position was removed from the average calculation. Error bars were determined by bootstrapping, with stratified sampling with replacement read coverage from individual PREs in each group separately.

### 4.6. Determination of PRE clustering

To determine whether the PREs cluster together more than would be expected by chance, we determined the ratio of the observed frequency of PUM sites within all possible 100 bp windows of 3′ UTRs with a least 1 PRE in them to a Poisson model with the rate parameter, *λ*, set to the average count of PREs within all 100 bp windows. 95% confidence intervals were determined by bootstrapping the observed distribution of PRE counts within all windows.

### 4.7. Predicting PUM-mediated regulation using conditional random forest models

In order to predict the PUM-mediated regulation on a given transcript, we used conditional random forest models as implemented by the cforest function from the party R package [96–98]. Binary classification models were trained using default settings with no parameter tuning on the Bru-Seq EFFECT and NOEFFECT classes and a permutation-based AUC variable importance metric was calculated for each individual model [78]. Due to the large class imbalance, ten separate datasets were generated from the full dataset, where the majority NOEFFECT class was randomly downsampled to match the EFFECT class. Within each of the ten datasets, five-fold cross validation was performed to assess performance and detect overtraining. Final models were generated using the ten downsampled datasets without cross-validation and performance was tested on the RNA-seq dataset from Bohn et al. [34]. Precision-recall plots were calculated using the PRROC package based on the methodology of Davis and Goadrich [99].

#### 4.7.1. Calculation of features associated with a PWM

For each of the features described, the values were first calculated individually for each unique 3′ UTR isoform. Values for each isoform were combined by taking the mean of the value for that feature and isoform weighted by the number of isoforms that shared that unique 3′ UTR in the full set of annotated 3′ UTRs in the hg19 genome. For features ending in “fimo_best_bygene_max_fimo”, the maximum FIMO match score for each unique 3′ UTR isoform for that PWM was calculated by setting the p-value cutoff threshold in FIMO to 1.1, thereby allowing FIMO to consider every possible match for a given sequence. The maximum match score for each sequence was reported for each unique 3′ UTR isoform. For features ending in “fimo_best_bygene_total_num”, the total number of matching sites for a given unique 3′ UTR isoform was calculated as described above in the “Determination of matching PREs” section. For each sequence, the geometric average of FIMO scores for each matching PRE was calculated and reported in the “fimo_bygene_geom_avg_score”. The maximum match score, geometric average match score, and total match number was calculated for the SEQRS PUM1 round 5 PWM, SEQRS PUM2 round 5 PWM, Hafner et al. [37] PUM2 PWM, and each of the PWMs for human RBPs found in the CISBP-RNA database [100].

For PREs, the shortest distance to the 3′ UTR for any given PRE is converted to normalized coordinates (i.e., 0.0 is the 5′ end and 1.0 is the 3′ end) and reported in the “fimo_best_bygene_dist_3′’. For “fimo_bygene_at_content” the largest percentage AT content in a 100 bp window surrounding any PRE within a given sequence was reported. Similarly for “fimo_bygene_max_cluster”, the maximum number of full PRE sites within a sliding of 100 bp was calculated. For both of these features, windows were truncated at the 3′ and 5′ ends of the sequence.

Predicted miRNA sites were determined using default predictions (conserved sites of conserved miRNA families) from TargetScan release 7.2 [67]. Overlaps with PREs were calculated by counting miRNA sites within a 100 bp window surrounding each PRE. For 3′ UTRs with more than one PRE, the PRE with the maximum number of overlapping miRNA sites was considered.

#### 4.7.2. Calculation of in silico basepairing probabilities for PREs

For each identified PRE, the probability of the given PRE being base-paired within predicted secondary structure was calculated using RNAfold [101] by calculating the ensemble free energy of an unconstrained sequence *F*_*u*_ of 50 bp flanking each side of a given PRE and the ensemble free energy of a constrained sequence where no base within the PRE is allowed to form a base pair *F*_*c*_. The probability of the PRE being constrained from base-pairing can be calculated using:

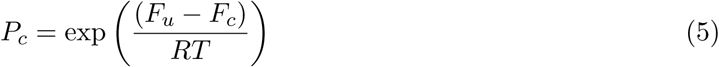

Where *T* is the temperature (set to physiological temperature, 310.15K), and *R* is the gas constant (set to 0.00198 kcal K^−1^ mol^−1^). Thus the probability of any given PRE being unpaired is *P*_*c*_. We define two features associated with *P*_*c*_ for each PRE in a given 3′ UTR isoform. “_avgprob_unpaired” is the average *P*_*c*_ of all the PREs within a given 3′ UTR and “_maxprob_unpaired” is the maximum *P*_*c*_ of all the PREs within a given 3′ UTR. Values for each isoform were combined into gene level estimates, as described above.

#### 4.7.3. Calculation of information redundancy between features

In order to calculate the information redundancy between features, each feature was discretized into ten equally populated bins. The redundancy between feature 1 (*F*_1_) and feature 2 (*F*_2_) was calculated with the following equation:

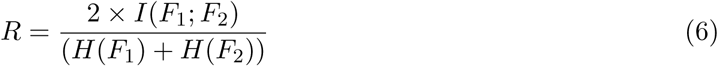

Where *H* is the entropy of a given vector *X* of discrete values, as defined below:

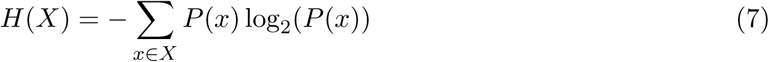

And the mutual information *I*(*X*; *Y*) of vectors *X* and *Y* of discrete values is defined as:

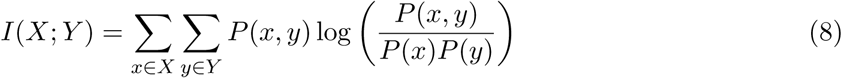

#### 4.7.4. Determination of EFFECT and NOEFFECT classes for RNA-seq data

RNA-seq data was obtained from Bohn et al. [34] and a gene was only considered if the FPKM for both the PUM1/2 knockdown condition and the siNTC condition were greater than 5. Genes that passed this cutoff and that were considered to have statistically significant differential expression in the original analysis were considered EFFECT genes. Genes that passed the cutoff and were not considered to have statistically significant differential expression were considered NOEFFECT genes.

## 5. Acknowledgments

This work was supported in part by the National Institute of General Medical Sciences, National Institutes of Health grant R35 GM128637 to P.L.F. and grant R01 GM105707 to A.C.G.. Additionally, this work was supported by National Institute of Neurological Disorders and Stroke, Grant/Award Number: R01NS100788 and NIH Grant/Award Number: 1UG3TR003149 to Z.T.C. as well as NIH Grant/Award Number: UM1 HG009382 and NIH Grant/Award Number: R01 CA213214 01 to M.L.. Work by M.B.W. was supported by the National Science Foundation Graduate Research Fellowship DGE1256260. Work by B.M. was supported by the NCI through the Rogel Cancer Center support grant P30CA046592.

## 5.1. Author contributions

Bioinformatics/computational analysis: Michael Wolfe (Data analysis and primary manuscript author), Peter Freddolino (Funding and concept, data analysis, writing); Bru-seq and BruChase-seq: Trista Schagat (RNAi, RNA labeling and purification), Aaron Goldstrohm (Funding and Concept, writing), Michelle Paulsen (BrU RNA Seq), Brian Magnuson (initial Bru-seq and BruChase-seq data analysis), Mats Ljungman (Funding and concept); PUM Protein Purification for SEQRS: Daeyoon Park, Chi Zhang and Zak Campbell (SEQRS and data analysis, funding)

---

**Figure S1:**
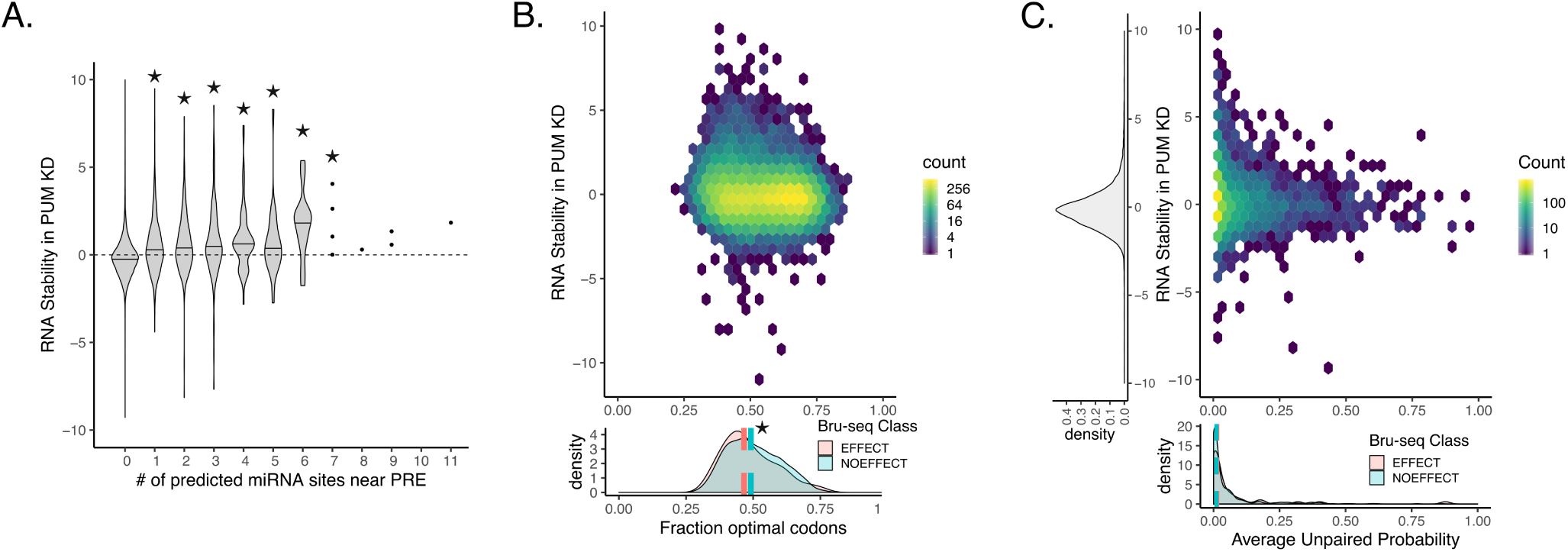
Additional features considered in determining PUM-mediated decay. A) Count of predicted conserved miRNA sites from conserved families that overlapped within 100 bp of a PRE for each gene. Stars indicate statistical significance from a Wilcoxon rank sum test compared to the 0 overlapping miRNA case. Stability in PUM knock-down is represented by a normalized interaction term between time and condition, where positive values indicate stabilization upon PUM knockdown and negative values indicate destabilization upon PUM knockdown (see Methods for details).B) (Above) Relationship between the fraction optimal codons as determined by the Codon Stability Coefficient determined in HEK293 cells [87] and PUM-mediated effect as measured in our Bru-seq data. (Below) Marginal density plots of the fraction optimal codons for genes in the EFFECT and NOEFFECT classes. Median fraction optimal codons for each class are plotted with dotted lines. A significant (p < 0.05, two-sided permutation test, n = 1000) difference in medians between the classes is indicated by a star. C) (Above) Relationship between the probability of a given PRE being unpaired in predicted RNA secondary structure. Only genes with a PRE with > 0 probability of being unpaired where shown in the heatmap. All other genes are shown in the marginal y-axis density plot. (Below) Marginal density plot for genes in the EFFECT and NOEFFECT classes with median probabilities for each class shown as dotted lines. See Methods for details of secondary structure prediction.

**Figure S2:**
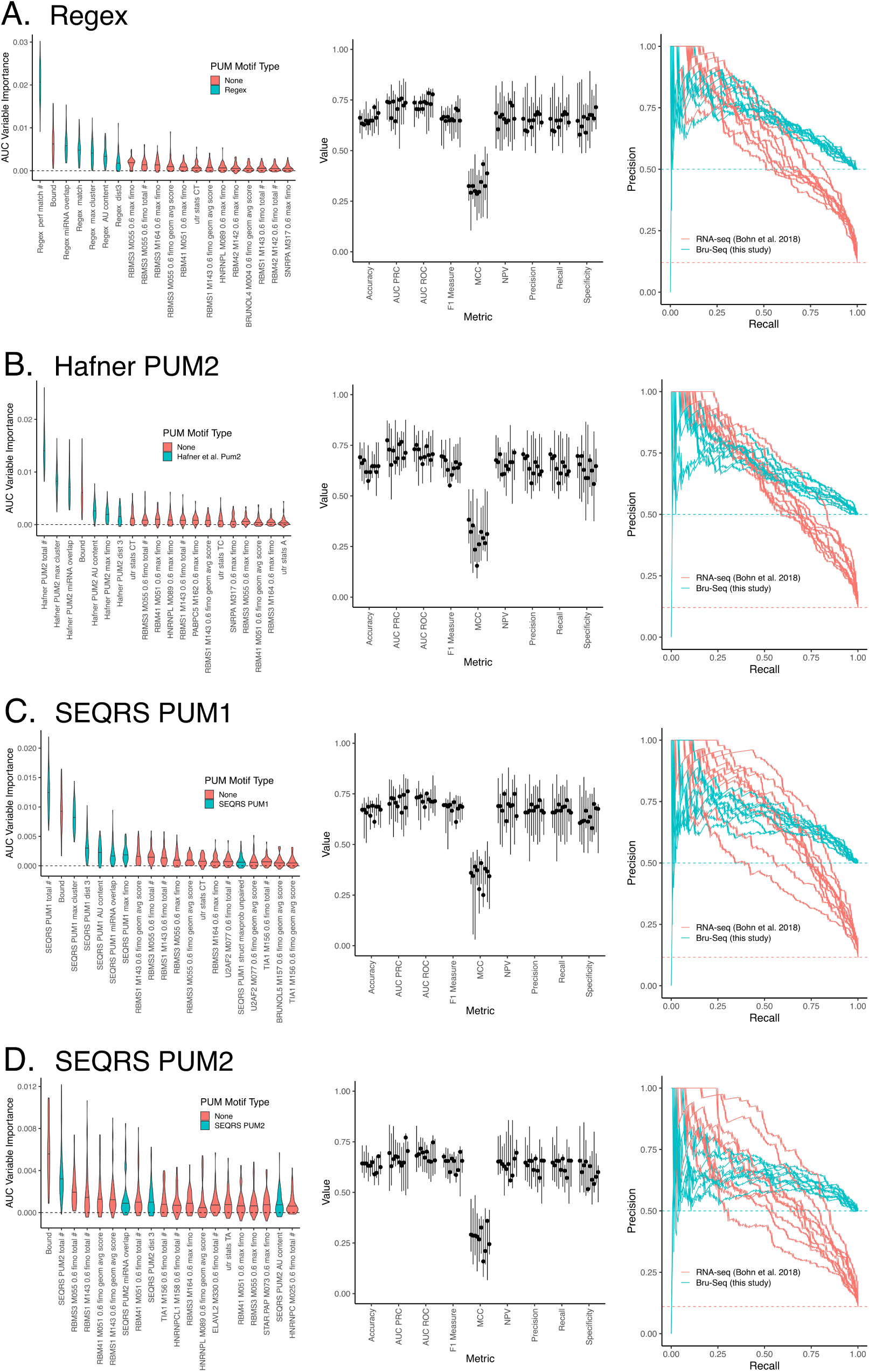
Predicting PUM-mediated effect subset by motif. A) Conditional random forest models for the datasets considering only genes that had at least one match to the regex motif definition in a 3′ UTR. PRE features only consider those around the regex definition. Panels are as in Figure 6B, D, and F. B) As in A), but for the Hafner et al. [37] PUM2 motif. C) As in A), but for the SEQRS PUM1 motif. D) As in A), but for the SEQRS PUM2 motif.

